# Different α_2_δ Accessory Subunits Regulate Distinctly Different Aspects of Calcium Channel Function in the Same Drosophila Neurons

**DOI:** 10.1101/838565

**Authors:** Laurin Heinrich, Stefanie Ryglewski

## Abstract

Voltage gated calcium channels (VGCCs) regulate neuronal excitability and translate activity into calcium dependent intracellular signaling. The pore forming α_1_ subunit of high voltage activated (HVA) VGCCs operates not in isolation but associates with α_2_δ accessory subunits. α_2_δ subunits can affect calcium channel biophysical properties, surfacing, localization and transport, but their *in vivo* functions are incompletely understood. In vertebrates, it is largely unknown whether different combinations of the four α_2_δ and the 7 α_1_ subunits mediate different or partially redundant functions or whether different α_1_/α_2_δ combinations regulate different aspects of VGCC function. This study capitalizes on the relatively simpler situation in the Drosophila genetic model that contains only two genes for HVA calcium channels, one Ca_v_1 homolog and one Ca_v_2 homolog, both with well-described functions in different compartments of identified motoneurons. We find that both dα_2_δ_1_ and dα_2_δ_3_ (stj) are broadly but differently expressed in the nervous system. Both are expressed in motoneurons, but with differential subcellular localization. Functional analysis reveals that dα_2_δ_3_ is required for normal Ca_v_1 and Ca_v_2 current amplitudes and for correct Ca_v_2 channel function in all neuronal compartments, axon terminal, axon, and somatodendritic domain. By contrast, dα_2_δ_1_ does not affect Ca_v_1 or Ca_v_2 current amplitudes or presynaptic function, but it is required for correct Ca_v_2 channel allocation to the axonal versus the dendritic domain. Therefore, different α_2_δ subunits are required in the same neurons to precisely regulate distinctly different functions of HVA calcium channels, which is in accord with specific α_2_δ mutations causing different brain diseases.

**Significance Statement:** Calcium influx through the pore forming α_1_-subunit of voltage gated calcium channels serves essential neuronal functions, such as synaptic vesicle release, control of action potential shape and frequencies, synaptic input computations, and transcriptional control. Localization and function of α_1_-calcium channel subunits depend on interactions with α_2_δ accessory subunits. Here we present *in vivo* analysis of Drosophila motoneurons revealing that different α_2_δ subunits independently regulate distinctly different aspects of calcium channel function in the same neuron, such as current amplitude and dendritic versus axonal channel localization. Our findings start unraveling how different α_1_/α_2_δ combinations regulate functional calcium channel diversity in different sub-neuronal compartments, and may provide an entry point toward understanding how mutations of different α_2_δ genes underlie brain diseases.

## Introduction

α_2_δ-accessory subunits affect multiple aspects of neuronal high voltage activated calcium channel (HVA VGCC) function (Brockhaus et al., 2018; Ferron et al., 2018; Tong et al., 2017; Fell et al., 2016; Hoppa et al., 2012). Consequently, mutations in α_2_δ genes are cause to neurological conditions such as ataxia (Calandre et al., 2016; Davies et al., 2007; Klugbauer et al., 2003), epilepsy (Faria et al., 2017; Celli et al., 2017; Dolphin, 2012; 2013), and neuropathic pain (Chen et al., 2019; Nieto-Rostro et al., 2018, Bauer et al., 2009). However, despite numerous reports on the roles of α_2_δ subunits in HVA channel trafficking (Kadurin et al., 2017; Hendrich et al., 2008), surfacing (D’Arco et al., 2015; Cassidy et al. 2014), and biophysical properties (Savalli et al., 2016; Davies et al., 2010, Felix et al., 1997, Hobom et al., 2000, Cantí et al., 2005), the specific *in vivo* functions that may result from different α_2_δ/α_1_ combinations remain incompletely understood. In heterologous expression systems, full calcium current amplitudes require co-expression of α_2_δ and β with the pore forming HVA VGCC α_1_ subunit (Barclay et al., 2001; Brodbeck et al., 2002; Davies et al., 2006; 2010; Cantí et al., 2005; Hoppa et al., 2012; for review see Dolphin, 2013), largely independent of which α_2_δ subunit is used, indicating redundant functions of α_2_δ subunits. By contrast, mutations of single α_2_δ subunit genes cause brain disease, rendering functional redundancy unlikely to occur *in vivo*. In fact, different α_2_δ subunits show differential spatial expression in the vertebrate brain (Scott and Kammermeier, 2017; Nieto-Rostro et al., 2014; Cole et al., 2005). A fundamental question thus is whether different combinations of α_2_δ and α_1_ subunits mediate distinctly different functions in neurons, what these are *in vivo*, and whether a combinatorial code of different α_2_δ/α_1_/β combinations regulates subcellular aspects of HVA channel function *in vivo*.

The 7 vertebrate HVA VGCC genes (Catterall, 2011) comprise two families, four Ca_v_1 channels (Ca_v_1.1-Ca_v_1.4) and three Ca_v_2 channels (Ca_v_2.1-Ca_v_2.3). In combination with four genes each for β- and α_2_δ-subunits (reviewed in Buraei and Yang, 2010; Dolphin, 2013), this totals to > 100 possible combinations of α_1_/β/α_2_δ HVA VGCC complexes. All so far tested α_1_/β/α_2_δ combinations are functional in heterologous expression systems, but it remains unclear how many of these are used *in vivo* to regulate different neuronal functions. To address this question we employ specific advantages of the Drosophila genetic model, which contains only one gene each homologous to the vertebrate Ca_v_1 and Ca_v_2 channel families. Dmca1D is homologous to the entire Ca_v_1 family, and Dmca1A, also named cacophony, to the Ca_v_2 family (Littleton and Ganetzky, 2000). Together with one β- and four genes encoding α_2_δ-subunits, this results in 8 possible combinations. Moreover, the relative simplicity, experimental accessibility, and available genetic tools make Drosophila a suitable system to study which α_2_δ/α_1_ combinations regulate which aspects of neuronal HVA channel functional diversity *in vivo*. We restricted the analysis to dα_2_δ_1_ and dα_2_δ_3_ subunits because high throughput expression data reveals high expression in the CNS (flybase.org). By contrast, dα_2_δ_2_ is reported muscle specific (Reuveny et al., 2018), and a predicted dα_2_δ_4_ lacks the double cache domain and shows little mRNA expression in the CNS.

We find differential expression of both α_2_δ subunits throughout the nervous system and during development. Surprisingly, both are expressed in larval and adult motoneurons, but localize to different subcellular compartments and mediate distinctly different functions. dα_2_δ_3_ is required for normal somatodendritic Ca_v_1 and Ca_v_2 current amplitudes, as well as for correct axonal and presynaptic Ca_v_2 channel function. By contrast, dα_2_δ_1_ is not required for presynaptic function or normal HVA current amplitudes but is critical for correct Ca_v_2 channel allocation to the axonal versus the dendritic domain. Thus different dα_2_δ subunits are required in the same neuron to cooperatively regulate distinctly different aspects of HVA channel function and localization.

## Materials and Methods

*Drosophila melanogaster* were reared at 25°C, on a 12/12hrs light/dark cycle, in plastic vials on a cornmeal, glucose, yeast, agar diet (for 6 liters: 725.69g glucose, 343.06g cornmeal, 66g Agar and 181.38g active dry yeast; after cooling to 70°C 76.25 ml Tegosept (10% in 100% ethanol) were added. After cooling to 65°C 3.5g ascorbic acid were added). Food was covered and left for 1 day at 4°C to cool and harden.

Experiments were carried out either in 2-5 day-old adult male and female, pupal stage P8 (Bainbridge and Bownes, 1981), and third instar larval animals of varying genotypes (for full list of genotypes see tables 1 and 2).

The Drosophila gene CG4587 is predicted to encode an α_2_δ subunit (Flybase) but until now no functional data existed. CG4587 was listed as the first dα_2_δ subunit (Kurshan et al., 2009). We used this list to name CG4587 dα_2_δ_1_. However, we do not claim that dα_2_δ_1_ fully resembles vertebrate α_2_δ_1_.

### Climbing Assay

2-4 day old single male or female virgins were put in separate plastic vials one day before testing. The climbing behavior was filmed, while a ruler was placed beside the vial as a measure of length. Gently hitting the vial on the ground will induce upwards climbing behavior due to negative geotaxis. Since flies tend to be more inactive during midday, testing was always done between 9:00am-1:00pm or 3:00-5:00pm at 25 °C. The climbing speed was analyzed manually with the Avidemux software. Length and duration of climbing attempts were measured to estimate the climbing speed. Mean climbing speed was calculated from three climbing events per fly.

### Immunocytochemistry

Immunohistochemistry for stj^mCherry^ and dα_2_δ_1_^GFP^ was conducted using antibodies against fluorescent epitopes that were fused to stj and dα_2_δ_1_ and genomically expressed because antibodies against Drosophila α_2_δ subunits are not commercially available.

(1) For triple staining of α-GFP, α-mCherry and α-Brp adult flies and larvae were dissected and fixated with 4% paraformaldehyde (PFA) in 0.1 M PBS (adults: 45 min / larvae: 30 min). Preparations were washed 3 times with PBS for 20 min and then permeabilized and blocked with 10 % normal donkey serum (NDS) in PBS-TritonX (0.5 %) for 5×60 min. Primary antibodies (chicken anti-GFP, 1:400, Invitrogen, A10262, RRID:AB_2534023; mouse anti-Brp, 1:200, nc82, DSHB, RRID:AB_2314867; rat anti-mCherry,1:5000, 16D7, cat# M11217) were diluted in 10% NDS in PBS-TritonX (0.3 %) and incubated at 4 °C overnight, shaking. Afterwards the preparations were washed with PBS 8 x 30 min and then incubated with secondary antibodies (donkey anti-chicken Cy2, 1:400, Dianova, cat# 703-505-155; donkey anti-rat Cy3, 1:400, Invitrogen, A-21209; donkey anti-mouse Cy5, 1:400, Invitrogen, A-31357) in PBS at 4 °C overnight, shaking. After washing the preparations 6 x 30 min in PBS, they were dehydrated with an ascending series of 50%, 70%, 90% and 100% ethanol for 10 min each and mounted in methylsalicylate.

(2) The staining of α-HA was done as described in (1). Instead of NDS, 5% bovine serum albumin (BSA) in PBS-TritonX (0.5 %) solution was used for blocking and permeabilization of the preparations. The primary antibodies (rat anti-HA, 1:100, Roche life science, 11867423001) were diluted in 5% BSA + PBS-TritonX (0.3 %). The secondary antibodies (donkey anti-rat Cy5, 1:400, Jackson Immuno Research, 712-605-150) were diluted in PBS.

### Quantification of axonal cac^GFP^ label

For live detection of axonal cacophony^GFP^ (cac; Ca_v_2^GFP^) label, a genomically tagged and endogenously expressed cacophony carrying an N-terminal GFP-tag was used (Gratz et al., 2019).

For quantification of axonal cac^GFP^ label all preparations were treated exactly the same way. 2-3 day old flies were dissected and instantly fixated with ice-cold 100% ethanol for 10 min. After washing the preparation with PBS for 10 min it was mounted in Vectashield (Vector Laboratories, Lot# X1215) and directly scanned with a Leica TSC SP8 Laser Scanning Microscope (Leica Microsystems Inc., RRID:SSR_004098) with excitation wavelength at 488 nm (Argon laser). All samples were scanned with a 20x glycine objective, a zoom factor of 1.8 for further magnification, a z-step size of 0.3 µm and an image resolution of 1024×1024 pixels. Furthermore, laser and detector settings were always identical and the laser was always warmed up 1 h before images were taken.

Projection views of the axons from stacks of 10 sections were analyzed with Fiji ImageJ 64 V5. To calculate the intensity of the axonal cac^GFP^ label a section of the axon, shortly after leaving the ventral nerve cord, was encircled and the mean gray value was measured. Per fly, the mean axonal cac^GFP^ intensity was calculated from the axons of both sides.

### Western Blotting

L3 larvae were stunned on ice for 5 min and dissected in ice-cold saline. Afterwards the CNS (α_2_δ_1_^GFP^: 20; stj^mCherry^: 30) were collected in 70 µl ice-cold 2xSDS sample buffer (25ml 4x Tris CI/SDS pH 6.8, 20ml glycerol, 4g SDS, 0.31g DTT, 1 mg Bromophenol Blue, add to 100ml with ddH_2_0), homogenized and boiled at 96 °C for 3 min. Samples were then stored at −28 °C.

Discontinuous SDS-PAGE in a large Hoeffer gel chamber with 1.5 mm thickness and 15 pockets with 100 µl volume each was done. A 5 % (bis-acrylamide) stacking gel (6.8 ml ddH_2_0, 1.7 ml 30% bis-acrylamide, 1.25 ml 4x Tris/SDS pH 6.8, 100 µl 10% ammonium persulfate, 10 µl TEMED) and an 8% (bis-acrylamide) running gel (18.6 ml ddH_2_0, 10.7 ml 30% bis-acrylamide, 10 ml 4xTris/SDS pH 8.8, 400 µl 10% ammonium persulfate, 16 µl TEMED) was poured and polymerized at 37 °C. Afterwards the pockets were washed with SDS-glycine-Tris electrophoresis buffer (3 g Tris base, 14.4 g glycine, 1 g SDS, add to 200ml with ddH_2_O). Samples were again boiled at 96 °C and centrifuged at 10,000 g for 1 min before loading. As a marker 70 µl of Color Protein Standard Broad Range (New England BioLabs, #P7712S; 25 to 245 kD) diluted 1/7 in SDS sample buffer was loaded. The gel was run at 0.02 A until the dye front passed the stacking gel, then the current was increased to 0.03 A (PowerPac, Bio-Rad).

Proteins were blotted onto nitrocellulose in a large wet tank filled with transfer buffer (18.2g Tris base, 86.5g glycine, 900 ml methanol add ddH_2_O to 6L). The blotting of the proteins was done at 4 °C overnight at 40 V (PowerPac, Bio-Rad).

After blotting, the membrane was cut in half at about 80 kDa. The two membrane pieces were washed with ddH_2_O for 10 min, incubated with TBST (10 ml 1M Tris pH 7.5, 30 ml 5 M NaCl, 1 ml Tween20 add to 1000 ml with ddH_2_O) 3 times for 20 min and blocked with 10% dried milk-TBST solution or BlockAce-TBST solution (BlockAce, Bio-Rad, #170223) for 2 h. After washing the membrane pieces in TBST for 3 times 20 min, they were incubated separately with primary antibody (245-80 kDa: rabbit anti-GFP, 1:1,000, Life Technologies, A11122 / rabbit anti-mCherry, 1:1,000, Abcam [EPR20579]; 80-25 kDa: mouse anti-actin, 1:10,000, dshb JLA20) diluted in 2,5 % milk-TBST or 25% BlockAce-TBST solution at 4 °C overnight. Both membrane pieces were then separately washed with TBST 3 times for 20 min before incubation with secondary antibodies (245-80 kDa: goat anti-rabbit IgG, 1:10,000, Jackson ImmunoResearch, cat# 111-035-144; 80-25 kDa: goat anti-mouse IgG, 1:4,000, Millipore, 12-349) diluted in TBST for 2 h at 25 °C. After washing the membrane pieces 3 times for 20 min with TBST and 20 min with TBS membrane was incubated in Immobilon Western chemiluminescent HRP substrate (Millipore, cat# WBKLS0500) for 5 min. Bands were detected with a Fusion SL Camera and Fusion software (Vilber Lourmat). For analysis a profile blot of the western was done with Fiji ImageJ V5 and the integrated areas of the bands of interest were measured. The relative densities were calculated by dividing the bands of interest with their respective loading control (actin).

### Generation of stj^mCherry^ flies

Flies with endogenously mCherry-tagged stj were generated, as already described, with a *Minos* mediated integration cassette (MiMIC) (Venken *et al*., 2011). A MiMIC cassette is flanked by two inverted ϕC31 bacteriophage attP sites and contains a gene-trap cassette and the yellow^+^ marker. ϕC31 expression was driven by the *vasa* promoter. Flies with a MiMIC construct in a coding intron of stj were obtained from Bloomington Drosophila Stock Center (BDSC_34109) and a for the splicing phase (phase 0) compatible plasmid containing the *mCherry* sequence was obtained from the Drosophila Genomics Resource Center (DGRC #1299_pBS-KA-attB1-2-PT-SA-SD-0-mCherry; Venken et al., 2011).

Female virgins of a *vasa* integrase line (BDSC_36312) were crossed with the stj-MiMIC flies (BDSC_34109). F1 Stage 2 embryos were injected with the DNA solution containing the mCherry plasmid (300-400 ng/µl). The injection electrodes (Science Products, GB100TF-8P) were pulled with a Flaming/Brown micropipette puller (Sutter Instruments Co., Model P-97) and broken individually. Injections were conducted with a Femtojet Injector (Eppendorf, cat# 5253000017) in Voltalef 10s oil. After injection, embryos were covered with Voltalef 3S oil and kept on 25 °C. Hatched larvae were then raised on instant fly food. Every hatched fly was crossed individually with either female virgins or males of a balancer stock (y^1^w*; Cyo/Sna^Sco^). F1 offspring displaying the yellow phenotype were re-crossed with balancer flies to build a stock. All stocks were checked for correct integration of the *mCherry* construct via PCR.

Primer sequences were obtained from Venken et al., 2011: Orientation-MiL-F: GCGTAAGCTACCTTAATCTCAAGAAGAG; Orientation-MiL-R: CGCGGCGTAATGTGATTTACTATCATAC; mCherry-Seq-F: ACGGCGAGTTCATCTACAAG; mCherry-Seq-R: TTCAGCCTCTGCTTGATCTC. Four different PCR reactions (2µl 10x Thermopol buffer, 0.5 µl 10mM dNTP’s, 0.5 µl F-primer, 0.5 µl R-primer, 0.1 µl Taq polymerase, 1 µl DNA, 5.4 µl ddH_2_O^RNAse free^) had to be performed for each event. The following primer combinations were used: (1) Orientation-MiL-F / mCherry-Seq-R; (2) Orientation-MiL-F / mCherry-Seq-F; (3) Orientation-MiL-R / mCherry-Seq-R; (4) Orientation-MiL-R / mCherry-Seq-F and a touchdown PCR (Biometra®, TGradient, Labexchange) was performed: 1x (94 °C, 600 s); 8x (94 °C, 30 s; 68 °C + −1 °C, 30 s; 68 °C 90 s); 32x (94 °C, 30 s; 60 °C, 30 s; 68 °C, 90 s); 1x(68 °C, 600 s). PCR products were loaded on a 0.7% agarose gel with ethidiumbromide added directly to the gel and run at 70 V (PowerPac, Bio-Rad) for about 60 min. Correct integration was marked by positive primer reactions for the primer combinations 1 & 4.

### Repairing the α_2_δ_1_^RNAi^ stock (Vienna Drosophila Resource Center; VDRC_108150)

As previously described (Green *et al*., 2014), during generation of VDRC “KK” RNAi stocks in rare cases the RNAi construct integrated into a second landing site (40D) in addition to the intended 30D landing site. Integration of the construct in both sites can lead to expression of a toxic protein called Tiptop (Tio). Thus, in order to prevent unspecific effects, the used KK stocks needed to be tested via PCR. Primer sequences were used as described (Green et al., 2014) (C_Genomic_F: GCCCACTGTCAGCTCTCAAC; NC_Genomic_F: GCTGGCGAACTGTCAATCAC; pKC26_R: TGTAAAACGACGGCCAGT; pKC43_R: TCGCTCGTTGCAGAATAGTCC). Four primer reactions (2µl 10x Thermopol buffer, 0.5 µl 10mM dNTP’s, 0.5 µl F-primer, 0.5 µl R-primer, 0.1 µl Taq polymerase, 1 µl DNA, 5.4 µl ddH_2_O^RNAse free^) had to be done for each tested line (1. C_Genomic_F / pKC26_R; 2. C_Genomic_F / pKC43_R; 3. NC_Genomic_F / pKC26_R; 4. NC_Genomic_F / pKC43_R) and a touchdown PCR was performed: 1x (95 °C, 120 s); 5x (95 °C, 15 s; 68 °C + −1 °C, 15 s; 72 °C 50 s); 29x (95 °C, 15 s; 62 °C, 15 s; 72 °C, 50 s); 1x (72 °C, 120 s). PCR products were loaded a 0.7% agarose gel with ethidiumbromide added directly to the gel and run at 70 V (PowerPac, Bio-Rad) for about 60 min.

Integration of the construct into the 40D landing site resulted in a PCR product of approx. 450 bp (C_Genomic_F / pKC26_R), while an empty site resulted in a PCR product of approx. 1050 bp (C_Genomic_F / pKC43_R). Integration of the construct into the 30D site resulted in a PCR product of approx. 600 bp (NC_Genomic_F / pKC26_R), while an empty site resulted in a PCR product of approx. 1200 bp (C_Genomic_F / pKC43_R).

For the *α*_*2*_*δ*_*1*_^*RNAi*^ line (VDRC_108150) the pKC26 vector indeed integrated into both the 30D and 40D site. The unwanted 40D insertion was removed via miotic recombination. Female *α*_*2*_*δ*_*1*_^*RNAi*^ virgins (VDRC_108150) were crossed to males in which both sites were empty (VDRC_60100). Female virgins of the F1 progeny were then crossed to a second chromosome balancer stock. Putatively recombinant offspring could be pre-selected via eye color (red eyes) and were tested for one-sided recombination via PCR as described above.

### *In situ* electrophysiology and calcium imaging experiments

Voltage clamp and current clamp experiments (Schützler et al., 2019; Kadas et al., 2017; Ryglewski et al., 2012; Ryglewski et al., 2014) and calcium imaging experiments were carried out as published (Ryglewski et al., 2017).

An upright Zeiss Axio Examiner A1 epi-fluoresecence microscope with a 40x water dipping lens (Zeiss, Germany) with a fixed stage (Narishige) was used. Recordings were done at room temperature (24°C). Electrophysiological experiments were conducted from crawling MN somata in third instar larvae, and wing depressor MN somata (DLM, specifically MN5) from pupae stage P8 (∼47-50 hrs after puparium formation, approx. halfway through pupal development (P50%)) and 2-5 day-old adult *Drosophila melanogaster* of each sex. Selection criterion for P8 was orange eyes as visible through the pupal case (Bainbridge and Bownes, 1981). All electrophysiological recordings were carried out in patch clamp whole cell configuration with an Axopatch 200B patch clamp amplifier (Molecular Devices), either in voltage clamp or current clamp mode. Data were digitized at a sampling rate of 50 kHz using a Digidata 1440 analog/digital converter (Molecular Devices) and low pass filtered with a 5 kHz Bessel filter. Data were acquired with pClamp 10.7 software (Molecular Devices).

The ganglionic sheath of the ventral nerve cord was focally digested and debris was carefully loosened and removed from the MN membrane with 1% *Streptomyces griseus* protease type XIV in saline using a broken patch pipette (Ryglewski et al., 2012b), and then rinsed thoroughly. Recording patch pipettes were pulled with a PC-10 vertical electrode puller (Narishige) from 1.5 mm outer and 1 mm inner diameter patch clamp glass capillaries without filament (WPI, #PG52151-4). Pipette resistance with Ca^2+^ current recording solutions was ∼3.5 MΩ for pupal and adult MN5, ∼4 MΩ for larval MNs, in action potential recording solutions ∼6 MΩ for pupal and adult MN5, and ∼6.5 MΩ for larval MNs. For solutions see below. Preparations were perfused with fresh saline (∼0.5 ml/min) throughout the course of the entire experiment.

### Recording solutions

Intracellular Ca^2+^ current recording solution (in mM): 140 CsCl, 0.5 CaCl, 2 Mg-ATP, 11 EGTA, 20 TEA-Br, 0.5 4-AP, 10 HEPES; pH was adjusted to 7.24 with 1 N CsOH, osmolality was 327 mOsM/kg.

Extracellular Ca^2+^ current recording solution (in mM): 93 NaCl, 5 KCl, 4 MgCl_2_, 1.8 CaCl_2_, 1.8 BaCl_2_, 30 TEA-Cl, 2 4-AP, 5 HEPES, ∼35 sucrose. pH was adjusted to 7.24 with 1N NaOH, osmolality was adjusted to 320 mOsM/kg with sucrose if necessary. TTX was added directly to the bath (the perfusion was halted for 5 min) at 10^−7^M (adults and pupae) or 4*10^−7^M (larvae) to block fast Na^+^ current. K^+^ channels were blocked with TEA and 4-AP.

Intracellular action potential recording solution (in mM): 140 K-gluconate, 2 Mg-ATP, 2 MgCl_2_, 11 EGTA, 10 HEPES. pH was adjusted to 7.24 with 1 N KOH, osmolality was adjusted to 300 mOsM/kg with glucose if necessary.

Extracellular action potential recording solution (normal saline; in mM): 128 NaCl, 2 KCl, 4 MgCl_2_, 1.8 CaCl_2_, 5 HEPES, ∼35 sucrose, pH was adjusted to 7.24 with 1 N NaOH, osmolality was adjusted to 290 mOsM/kg with sucrose if necessary.

Intracellular Ca^2+^ imaging solution (in mM): 140 K-gluconate, 2 Mg-ATP, 2 MgCl_2_, 10 phosphocreatine di tris, 0.3 Na_2_GTP, 10 HEPES. EGTA was omitted because of the presence of GCaMP6s. pH was adjusted to 7.24 with 1 N KOH, osmolality was 313 mOsM/kg.

Extracellular action potential recording solution (normal saline; in mM): 115.8 NaCl, 2 KCl, 4 MgCl_2_, 5 CaCl_2_, 5 HEPES, ∼35 sucrose, pH was adjusted to 7.24 with 1 N NaOH, osmolality was adjusted to 305 mOsM/kg with sucrose if necessary.

### Voltage clamp and current clamp experiments (incl. Ca^2+^ imaging experiments)

**For voltage and current clamp recordings,** offset was nulled manually while approaching the cell, applying gentle positive pressure to the patch pipette to avoid dilution of the tip with extracellular solution. After gigaseal formation, mode was changed to patch configuration (or on-cell), and the cell was clamped to −30 mV (for Ca^2+^ current recordings) or −70 mV (for AP recordings), respectively. Fast capacitance artifacts of the recording electrode were zeroed using the C-slow and C-fast dials of the amplifier, lag was 2 µs. Break-in was achieved by short and quick, gentle suction. Configuration was changed to whole cell, and cell capacitance as well as series resistance were compensated for using the whole cell cap and serial resistance dials of the amplifier. Only recordings with series resistances below 10 MΩ were continued. Usually, series resistance was ∼8 MΩ. Prediction was set to ∼98%, and compensation was between 40 and 50%. In Ca^2+^ current experiments, the cell was manually clamped to −70 mV in 20 mV increments, once all parameters were compensated. This was necessary because the Goldman potential with the given solutions was around 0 mV and therefore far away from the intended holding potential of −70 mV. Clamping the cell to −70 mV immediately often results in rupture.

#### Ca^2+^ current recordings

Ca^2+^ currents were recorded in voltage clamp mode. Currents were evoked by 200 ms voltage steps from −90 mV to +20 mV (adult and pupal MNs) or 0 mV (larval MNs) from a holding potential of −90 mV in 10 mV increments. Linear leak was calculated from the first three voltage steps and subtracted offline. Adult Ca^2+^ current consists of low (LVA) and high voltage activated (HVA) currents. The fast LVA was isolated by addition of the off-artifact to the on-artifact. LVA can only be observed in isolation between −70 and −40 mV. HVA activates around −30 mV and is also carried by cacophony which makes selective block of one component impossible (Ryglewski et al., 2012).

#### AP recordings

After break-in, parameters were adjusted as for Ca^2+^ current recordings (s.a.) to get an idea how healthy the cell is. Then we switched to current clamp mode. Only cells with a membrane potential ≤ −55 mV were used. Pupal (P8) action potentials (Ryglewski et al., 2014) were elicited by depolarizing ramp or square current injection. For Ca^2+^ imaging experiments, a 400 ms 1 nA max. amplitude ramp current injection was performed which reliably elicited a train of action potentials.

#### Ca^2+^ imaging

APs were elicited as described above, and the resulting changes in GCaMP6s (Chen et al., 2013) fluorescence were recorded and analyzed. An Orca Flash 4.0 LT CMOS camera (C11440-42U; Hamamatsu Photonics K.K.) with HOKAWO 3.10 software was used for image acquisition. Exposure time was 75 ms. Image series were streamed. Raw data were exported to MS Excel, and ΔF/F was calculated by [F(firing)-F(rest)]/F(rest) (Ryglewski et al., 2017). Regions of interest (ROI) were chosen in dendrites and axon.

### Intracellular muscle recordings from L3 larvae

EPSPs were recorded in HL3.1 saline with 0.5 mM Ca^2+^ (62.5 mM NaCl, 10 mM MgCl_2_, 5 mM KCl, 0.5 mM CaCl_2_, 10 mM NaHCO_3_, 5 mM Trehalose, 5 mM HEPES, 35 mM Sucrose; pH 7.24-7.25, osmolality 300-310 mOsM/kg; Feng et al., 2004). Electrodes were pulled from borosilicate glass capillaries (WPI, 1B100F-4) with a Flaming/Brown micropipette puller (Sutter Instruments Co., Model P-97). L3 larvae were dissected and the CNS was removed at the end of the dissection procedure by cutting the nerves as close to the CNS as possible. A sharp electrode (tip resistance 30 MΩ) filled with 3 M KCl was placed close to muscle M10 of a thoracic segment. As reference, a chlorinated silver wire was placed inside the bath solution. Offset and capacitance of the electrode were adjusted manually before the electrode was inserted into the muscle. Signals were amplified with an Axoclamp 2B intracellular amplifier in Bridge mode, digitized with a Digidata 1440 and recorded with pClamp 10.7 software (all Molecular Devices). Only data from muscles with a membrane potential of ≤ −50 mV were used for analysis. To evoke PSPs the respective nerve was sucked into and stimulated by a suction electrode (Sutter, BF100-50-10; broken individually). As reference, a thin silver wire wrapped around the suction electrode was used. Electrical stimuli with a duration of 0.5 ms and the minimal voltage needed (+1 V) for action potential generation were applied via an Isolated Pulse Stimulator (Model 2100, A-M Systems) and amplified by a Differential AC Amplifier (Model 1700, A-M Systems). A stimulus train of 0.5 Hz was applied for 20 s. EPSP amplitudes were analyzed with Clampfit 10.7. Per animal, the mean amplitude of 10 EPSPs was calculated.

#### Intracellular Dye Fill

Adult MN5 was filled as described previously (Ryglewski et al., 2017). Adult flies were dissected, and the ganglionic sheath was enzymatically digested. Then the very tip of a sharp glass microelectrode (borosilicate, outer diameter 1 mm, inner diameter 0.5 mm, with filament, Sutter BF100-50-10) pulled with a Sutter P97 Flaming Brown horizontal electrode puller was filled with a 50/50 mixture of TRITC-Dextran 3000 and Neurobiotin (both Thermo Fisher) in 2 M KAcetate. Then the shaft was filled with 2 M KAcetate leaving an air bubble between the dye-loaded tip and the KAcetate to avoid dye dilution. The electrode was connected to an intracellular amplifier (Axoclamp 2B, Molecular Devices) in Bridge mode; tip resistance was ∼60 MΩ. After impalement of the MN soma with the sharp electrode, the dye was injected iontophoretically into the cell by application of up to 1 nA positive current. Filling quality was judged visually. After completion, the electrode was removed, and the preparation was fixed with 4% paraformaldehyde in phosphate buffered saline (PBS) for 50 minutes at room temperature. After fixation the preparation was washed at least 6×20 min with PBS, then 6×20 min with 0.5% PBS-TritonX 100, both shaking. This was followed by incubation in Streptavidin coupled to Cy3 at a concentration of 1:750 at 4°C overnight, shaking. The preparation was then rinsed a few times with PBS, and then washed at least 6×30 min with PBS. Then the preparation was subjected to an ascending ethanol series (50, 70, 90, 100% ethanol), 10 min each, and then mounted in methylsalicylate on metal slides with a 8 mm whole with a glass cover slip glued to one side with super glue. Preparations were covered with a glass cover slip, which was sealed with nail polish. Reconstruction-ready images are generated using a Leica TSC SP8 confocal laser scanning microscope with a 40x, 1.25 NA oil lens with a 561 nm DPSS laser. Detection range was between 570 and 600 nm. Z-step size was 0.3 nm, zoom 3.5. Voxel dimensions were 86 x 86 x 290 nm (x, y, z). Dendritic structure was reconstructed from confocal image stacks after export to Amira software (AMIRA 4.1.1, FEI, Hillsboro, Oregon, US) with custom plug-ins (Schmitt et al., 2004; Evers et al., 2005).

## Results

### stj and dα_2_δ_1_ are differentially expressed through the CNS but co-localize in motoneurons

To assess the localization of dα_2_δ_1_ (CG4587) and dα_2_δ_3_ (straightjacket, stj) in the Drosophila ventral nerve cord (VNC), we used genomically tagged protein trap (Nagarkar-Jaiswal et al., 2015) fly strains (stj^mCherry^, dα_2_δ_1_^GFP^, see methods). In both, the larval (Figs. 1A-C) and the adult VNC (Figs. 1D-G) stj (green) and dα_2_δ_1_ (magenta) are broadly expressed, indicating functional relevance in many neurons. dα_2_δ_1_ localizes to all neuropil regions of the larval (Figs. 1Ai, Bi) and the adult VNC (Fig. 1Di, Ei), which comprise axon terminals and dendrites and are counter-labeled with the synaptic marker, bruchpilot (brp, cyan, Aii, Bii, Dii, Eii; Wagh et al., 2006; Kittel et al., 2006). By contrast, stj was not detected in central neuropils (Figs. 1B, D, E). This differential localization pattern was also found during pupal life (not shown) thus indicating different functions of stj and dα_2_δ_1_ at all stages of post-embryonic nervous system development. Localization of stj in the ganglionic cortex (somata) but not in neuropils was further confirmed by expression of an HA-tagged *UAS-stj-HA* transgene under the control of the *stj-GAL4* promoter (Figs. 1C, G). HA-label reveals stj expression in MN somata in the larval (Fig. 1C, asterisks) and in the adult VNC (Fig. 1G, arrows). However, we cannot exclude low expression levels below immunocytochemical detection levels of stj in neuropil regions. Importantly, despite their differential localization with respect to VNC synaptic neuropil and ganglionic cortex, both stj and dα_2_δ_1_ also co-express in many neuronal somata, including identified larval crawling MNs (Figs. 1A-Aiv, white arrows in Aiii, Aiv, and Merge) and adult wing depressor (DLM) MNs (Figs. 1E-Fi). Due to the abundant localization of dα_2_δ_1_ in the neuropil, expression in wing MN somata is difficult to see (Fig. 1Ei, arrow heads), but was confirmed by visual inspection of single optical sections and in pupal VNC with less dense neuropil (Figs. 1F, Fi, MN somata marked with arrow heads and dotted white circles).

**Figure 1:**
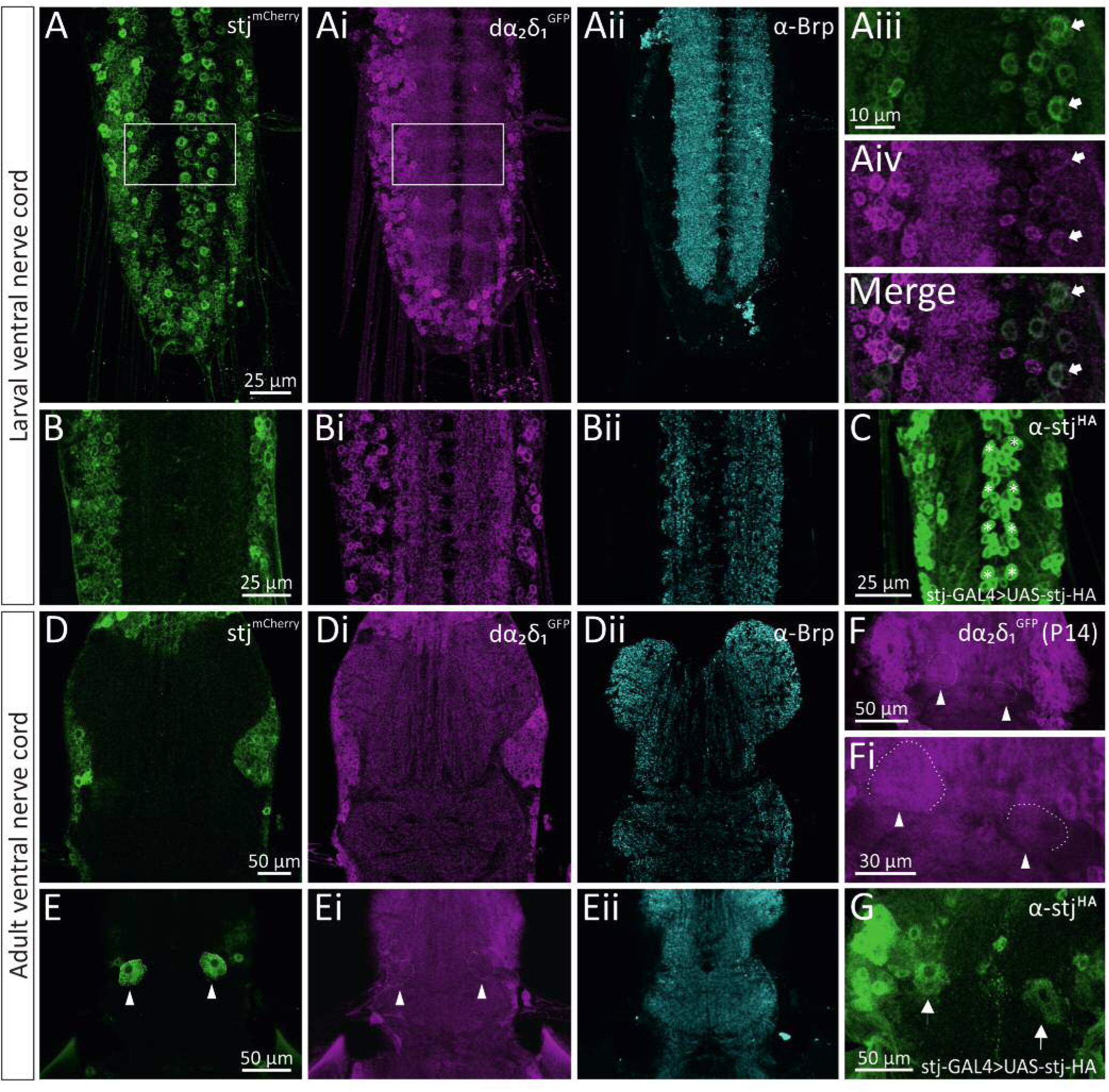
stj and dα_2_δ_1_ are differentially expressed in the Drosophila CNS but co-localize in motoneurons (MNs). (**A-Aii**) Projection view of 40 dorsal optical sections from a representative confocal image stack of the larval ventral nerve cord (VNC). **(A)** stj^mCherry^ (green) and **(Ai)** dα_2_δ_1_^GFP^ (magenta) are both expressed in many somata in the ganglionic cortex of the larval VNC, including overlapping expression in crawling MNs. **(Aii)** Neuropil is labeled with the synaptic marker brp (cyan). No stj^mCherry^ (**A**, green) but abundant dα_2_δ_1_^GFP^ (**Ai**, magenta) was found in the neuropil. **(Aiii, Aiv, Merge)** For better visualization of expression in MN somata five sections were selectively enlarged at the regions marked by white rectangles in (**A**) and (**Ai**). Expression of both stj^mCherry^ (**Aiii**) and dα_2_δ_1_^GFP^ (**Aiv**) in larval crawling MNs (white arrows). The (**Merge**) shows an overlay of (**Aiii**) and (**Aiv**) revealing co-label in MNs. (**B-Bii**) Same representative preparations as in (**A-Aii**), but deeper in the VNC and at higher magnification to visualize neuropil regions. **(B)** The neuropil is devoid of stj^mCherry^ (green), which is localized only to somata in the ganglionic cortex. **(Bi)** dα_2_δ_1_^GFP^ (magenta) is abundantly expressed in VNC synaptic neuropil that is counterstained with brp (**Bii**, cyan). (**C**) stj-HA is expressed under the control of the stj promoter (stj-GAL4>UAS-stj-HA) in the larval VNC. Neuropil label is not detectable, but stj-HA is found in many somata in the larval VNC including MNs (asterisks). **(D-E)** Adult VNC. **(D)** stj^mCherry^ (green) and dα_2_δ_1_^GFP^ (**Di**, magenta) are expressed in somata in the ganglionic cortex. **(Di, Dii)** dα_2_δ_1_^GFP^ is abundantly expressed in neuropil regions (**Di**, magenta) which are co-labeled with the synaptic marker brp (**Dii**, cyan). (**E**) Expression of stj (green) in adult wing depressor MNs (white arrowheads). **(Ei)** High expression levels of dα_2_δ_1_ (magenta) in neuropil (**Eii**, brp, cyan) results in low contrast for detection in adult wing MNs. Localization of DLM MN5 is indicated by dotted white circles and arrow heads **(E, Ei).** (**F, Fi**) Therefore, expression of dα_2_δ_1_^GFP^ in wing depressor MNs was also confirmed in pupal stage P14, shortly before adult eclosion (see dotted white line in (**F**) and in the enlargement in (**Fi**). (**G**) stj-HA expressed under the control of stj-GAL4 is detectable in adult DLM MNs (arrows) and in other somata.

To investigate the functional consequences of stj and dα_2_δ_1_ malfunction, we targeted stj and dα_2_δ_1_ RNAi to MNs. Knock down efficacy was determined by Western Blotting following pan-neural expression of either stj or dα_2_δ_1_ RNAi (Fig. 2A). Although transgene expression levels in MNs may differ from average pan-neural expression levels, this approach yields a reasonable estimate of knock down efficacy. In controls, two bands were detected at the expected sizes of α_2_δ and α_2_ alone (Ferron et al., 2018). Knock down was 64% on average for elav^C155^-GAL4>UAS-stj^RNAi^ (Fig. 2Ai, left two bars) and 98% on average for elav^C155^-GAL4>UAS-dα_2_δ_1_ ^RNAi^ (Fig. 2Ai, right two bars).

**Figure 2:**
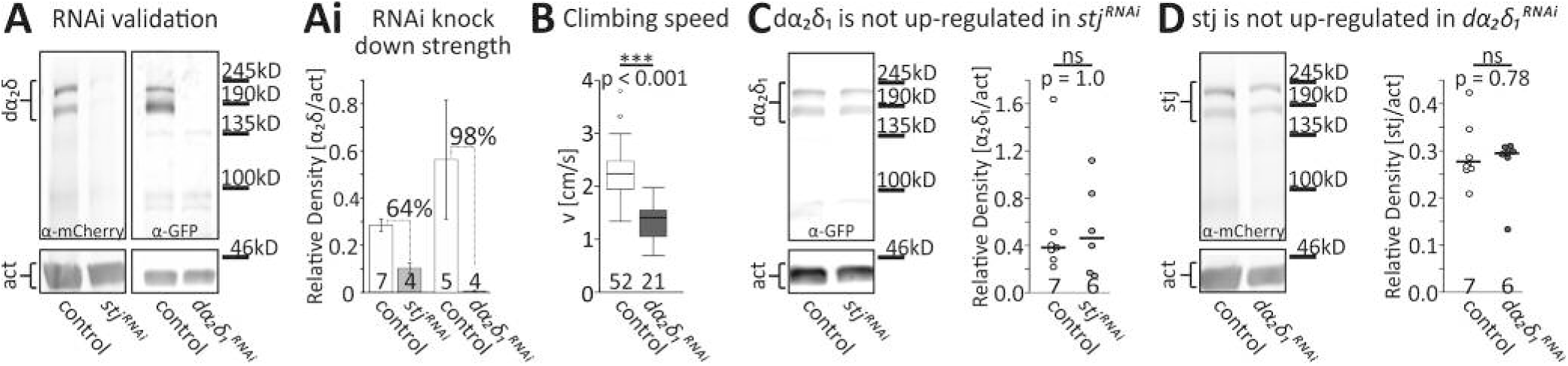
stj and dα2δ1 do not compensate for each other on the protein level. **(A, Ai)** Analysis of stj and dα_2_δ_1_ RNAi knock down efficacy. **(A)** Western blotting shows effective knock down of stj^mcherry^ (left two lanes) and of dα_2_δ_1_^GFP^ (right two lanes) following pan-neurally expressed *UAS-stj*^*RNAi*^ and *UAS-dα_2_δ*_1_^RNAi^ transgenes (*elav^c155^-GAL4>UAS-RNAi; UAS-dcr2*). The respective control lanes show two bands, because a fraction of α_2_δ is cleaved into α_2_ and δ. The upper α_2_δ band corresponds to un-cleaved α_2_δ, the lower band to cleaved α_2_ that carries the mCherry (stj) or GFP-tag (dα_2_δ_1_), respectively. **(Ai)** Quantification reveals 64% knock down efficacy for stj (control and knock down, left two bars), and 98% knock down on the protein level for *dα*_*2*_*δ*_*1*_ (right two bars). Numbers in bars indicate number of replicates. **(B)** *dα*_*2*_*δ*_*1*_^*RNAi*^ reduces climbing speed. In a negative geotaxis assay, pan-neural *dα*_*2*_*δ*_*1*_^*RNAi*^ results in a 40% median reduction in climbing speed from 2.3 to 1.4 cm/s as compared to control (N for control: 52, for *dα_2_δ_1_^RNAi^*: 21; p < 0.001, Mann-Whitney U-test). Boxes represent median and 25 and 75% quartiles. Whiskers represent 10 and 90% quartiles. **(C, D)** stj and dα_2_δ_1_ do not compensate for each other when either one is knocked down via pan-neural RNAi. *stj*^*RNAi*^ does not affect dα_2_δ_1_ protein level (**C**, p = 1, Mann-Whitney U-test), and vice versa, *dα*_*2*_*δ*_*1*_^*RNAi*^ does not affect stj protein level as analyzed by Western Blot (**D**, p = 0.78, Mann-Whitney U-test). Data in **(C)** and **(D)** are presented as single data point with median.

Both dα_2_δ_1_ and stj express in the same identified MNs, but for the following reasons they likely mediate different functions: First, stj localizes to somata (Fig. 1) and to neuromuscular terminals of MNs (Dickman et al., 2008; Ly et al., 2008; Kurshan et al,. 2009), whereas dα_2_δ_1_ is localized to somata and central neuropils. Second, stj loss of function is embryonic lethal but loss of dα_2_δ_1_ is viable, although it significantly reduces the speed of locomotion (Fig. 2B). In addition, *stj*^*RNAi*^ targeted to DLM MNs causes inability to fly, but *dα*_*2*_*δ*_*1*_^*RNAi*^ in the same MNs does not abolish flight. Finally, pan neural knock down of either dα_2_δ subunit did not result in compensatory upregulation of the other one as revealed by Western Blotting for stj following pan neuronal knock down of dα_2_δ_1_ and *vice versa* (Figs. 2C, D). Therefore, we hypothesize that both dα_2_δ subunits are required in MNs for different functions.

### stj but not dα_2_δ_1_ is required for normal MN Ca_v_1-like and Ca_v_2-like current amplitudes in vivo

To assess the functions of dα_2_δ in motoneurons we targeted RNAi knock down selectively to larval crawling MNs and recorded neuromuscular transmission (Figs. 3A, Ai) and somatodendritic Ca^2+^ currents (Figs. 3B-C). An ∼60% reduction of stj protein expression in MNs by targeted expression of *stj*^*RNAi*^ reduced larval neuromuscular transmission by ∼50%, as revealed by current clamp recordings of EPSPs from muscle 10 following extracellular stimulation of the motor nerve (Figs. 3A middle trace, 3Ai light gray box). By contrast, *dα*_*2*_*δ*_*1*_^*RNAi*^ did not reduce the amplitude of neuromuscular transmission (Figs. 3A, right trace, 3Ai, dark gray box), indicating that dα_2_δ_1_ is not required for normal Ca_v_2 channel function in MN axon terminals.

**Figure 3:**
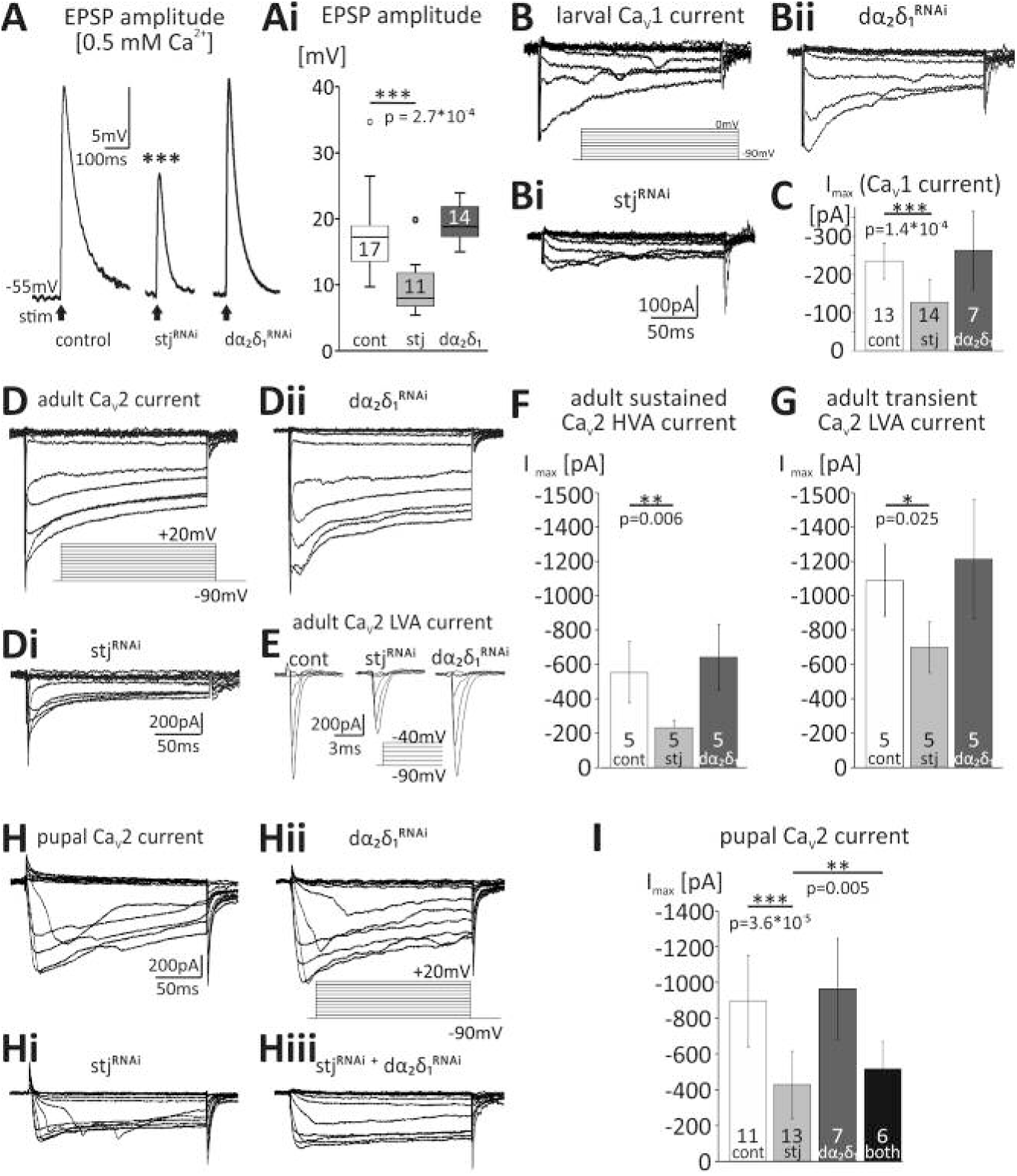
stj but not dα_2_δ_1_ is required for normal MN Ca_v_1-like and Ca_v_2-like current amplitudes in vivo. **(A, Ai)** Analysis of larval EPSPs. **(A)** Effect of *stj*^*RNAi*^ and of *α*_*2*_*δ*_*1*_^*RNAi*^ targeted only to MNs (*vGlut^OK371^-GAL4>UAS-RNAi; UAS-dcr2*) on larval neuromuscular synaptic transmission was tested in sharp electrode muscle recordings of muscle 10. EPSPs were monitored following extracellular MN stimulation. *stj*^*RNAi*^ (middle trace) but not *dα*_*2*_*δ*_*1*_^*RNAi*^ (right trace) caused a reduction in EPSP amplitude as compared to control (left trace). **(Ai)** Quantification revealed a ∼50% and highly significant reduction in EPSP amplitude by *stj*^*RNAi*^ (light gray box) as compared to control (white box), but *α*_*2*_*δ*_*1*_^*RNAi*^ had no effect (dark gray box). control n=17, *stj*^*RNAi*^ n=11, *α*_*2*_*δ*_*1*_^*RNAi*^ n=14; **p=0.001, Kruskal Wallis ANOVA with Dunn’s post hoc test). Boxes display median with 25 and 75% quartiles, whiskers represent 10 and 90% quartiles. **(B-I)** Whole cell Ca^2+^ current recorded from larval crawling (**B-C**), adult **(D-G)**, and pupal DLM MNs **(H-I)** *in situ*. Currents were elicited from a holding potential of −90 mV in 10 mV command voltage steps of 200 ms duration to 0 mV (larva) or +20 mV (adult and pupae) using blockers for Na^+^ and K^+^ channels. For voltage protocols see insets in **(B)**, (**D**) and (**Hii**). As compared to control **(B, C**, n=13) larval somatodendritic Ca_v_1-like current amplitude was reduced by 46% following *stj*^*RNAi*^ (**Bi, C**, n=14; ***p=1.4*10^−4^, one-way ANOVA with LSD post hoc test) but not by *dα*_*2*_*δ*_*1*_^*RNAi*^ (**Bii, C**, n=7). **(D-I)** Adult and pupal Ca^2+^ currents were recorded from DLM MN5. Knock down was driven specifically in DLM MNs by *23H06-GAL4>UAS-RNAi; UAS-dcr2*. **(D-G)** Adult Ca_v_2 Ca^2+^ current. As compared to control **(D**, n = 5) adult HVA Ca_v_2 current was reduced by ∼60% following *stj*^*RNAi*^ (**Di, F**, n=5, **p=0.006, one-way ANOVA with LSD post hoc test), but unaffected by *dα*_*2*_*δ*_*1*_^*RNAi*^ (**Dii, G**, n=5). **(E)** Adult low voltage activated (LVA) Ca_v_2 current (control, left trace) was reduced by 36% following *stj*^*RNAi*^ (**E**, middle trace, **G**, *p=0.025, one-way ANOVA with LSD post hoc test) but unaffected by dα_2_δ_1_^RNAi^ (**Dii, G**). **(H-I)** Pupal stage P8 (P50%) HVA Ca_v_2 current. Pupal Ca_v_2 current was reduced by 53% on average following *stj*^*RNAi*^ (**H, Hi, I**, white and light gray bars, control n=11; *stj*^*RNAi*^ n=13, ***p=3.6*10^−5^, one-way ANOVA with LSD post hoc test) but unaffected by *dα*_*2*_*δ*_*1*_^*RNAi*^ (**Hii, I**, dark gray bar, n=7). Double *stj*^*RNAi*^/*dα*_*2*_*δ*_*1*_^*RNAi*^ did not reveal additional effects and reduced Ca_v_2 current amplitude by ∼43% as compared to control (**Hiii, I**, black bar, n=6; **p = 0.005, one-way ANOVA with LSD post hoc test). Data in **(C)**, **(F)**, **(G)**, and **(I)** are represented as means ± SD.

To further test whether different dα_2_δ subunits regulate HVA currents selectively in different sub-neuronal compartments, or different HVA channels, or both, we next recorded Ca_v_1 and Ca_v_2 somatodendritic Ca^2+^ current by somatic voltage clamp recordings of identified MNs. Larval MNs express Ca_v_2 like channels in the axon terminal active zones (Gratz et al., 2019), but somatodendritic Ca^2+^ current is mediated by the Ca_v_1 homolog, Dmca1D (Littleton and Ganetzky, 2000; Worrell and Levine, 2008; Kadas et al., 2017; Schützler et al., 2019). By contrast, adult and pupal DLM MNs use the Ca_v_2 homolog, Dmca1A (cacophony, Littleton and Ganetzky 2000,) for both, axon terminal (Kawasaki et al., 2004) and somatodendritic Ca^2+^ current (Ryglewski et al., 2012; 2014).

Following *stj*^*RNAi*^, MN Ca^2+^ current amplitudes were decreased on average by 46% in larval crawling MNs (HVA Ca^2+^ current, Figs. 3B, Bi, C), by 59% (sustained HVA, Figs. 3D, Di, F) or 36% (transient LVA, Figs. 3E, G) in adult DLM MNs, and by 53% in pupal DLM MNs (HVA Ca^2+^ current, Figs. 3H, Hi, I), respectively. On the contrary, *dα*_*2*_*δ*_*1*_^*RNAi*^ did not affect somatodendritic Ca^2+^ current amplitudes, neither larval Ca_v_1-like nor adult or pupal Ca_v_2-like current (Figs. 3B-I). Moreover, following knock down of both stj and dα_2_δ_1_ in the same pupal DLM MNs Ca^2+^ current amplitudes reflect that of stj knock down data (Figs. Hiii, I). In summary, *stj*^*RNAi*^ impairs both pre-synaptic function as well as somatodendritic Ca^2+^ currents but dα_2_δ_1_^*RNAi*^ does not. Hence, stj seems important for normal Ca^2+^ current amplitudes independent of channel type and developmental stage. We next addressed the role of dα_2_δ_1_ for which no functional data exist up to date in Drosophila.

### stj and dα_2_δ_1_ have opposite effects on functional VGCC expression in the axon

In addition to the prominent role of HVA VGCCs at the pre-synapse for action potential (AP) triggered synaptic vesicle release, and known dendritic functions, axonal functions of HVA channels have been described in both, larval Drosophila MNs (Kadas et al., 2017) and developing adult Drosophila wing MNs (Ryglewski et al., 2014). To visualize axonal Ca_v_2 channels on the level of confocal microscopy, we used genomically GFP-tagged cacophony (Ca_v_2, Littleton and Ganetzky, 2000) channels, which have been reported to function and localize not significantly differently from native channels (Gratz et al., 2019). The arrangement of all 5 DLM wing MN axons into one axon bundle exiting the VNC towards the DLM wing depressor muscle allows visualization of GFP-tagged Ca_v_2 channels in MN axons by confocal microscopy (Fig. 4A). Axonal Ca_v_2^GFP^ channel label was visibly (Fig. 4A, middle panel) and statistically significantly decreased by targeted *stj*^*RNAi*^ knock down (Fig. 4B). By contrast, *dα*_*2*_*δ*_*1*_^*RNAi*^ causes increased Ca_v_2^GFP^ channel label in MN axons (Figs. 4A, bottom panel, 4B). Therefore, stj and dα_2_δ_1_ have opposite effects on axonal cacophony channel abundance in DLM MNs. To test whether this was caused by altered transport of cacophony channels, or by functional channels in the membrane we next recorded AP shape in current clamp mode. The DLM MN action potential is mainly carried by Na^+^, but it also contains a Ca^2+^-component during specific stages of pupal life (^48^). This Ca^2+^ component can be uncovered by bath application of the potent, ubiquitous, and irreversible VGCC blocker Cd^2+^ (500 µM) that reduces AP width in controls (Fig. 4C, upper panel, left two traces, arrow head). Following *stj*^*RNAi*^ AP width was smaller than in controls (Figs. 4C), and bath application of Cd^2+^ did not decrease AP width (Figs. 4C, D), indicating that the Ca^2+^ component was missing. These data are in agreement with a reduced expression of functional Ca^2+^ channels in MNs axons following *stj*^*RNAi*^. By contrast, the Ca^2+^ component was even more pronounced and the AP was broadened following *dα*_*2*_*δ*_*1*_^*RNAi*^ (Fig. 4C). AP shape was affected to an extent that the Ca^2+^ shoulder (Fig. 4C, top left trace, arrow head; ^48^) amounted to a double peak that was abolished by application of Cd^2+^ (Figs. 4C, top and bottom traces, D). Therefore, *dα*_*2*_*δ*_*1*_^*RNAi*^ likely increases the density of functional Ca_v_2 channels in MN axons. A role of dα_2_δ subunits on axonal Ca^2+^ influx and thus AP shape is further supported by AP recordings in high Ca^2+^ recording saline (5 mM) which results in even broader APs in controls and following *dα*_*2*_*δ*_*1*_^*RNAi*^ but not *stj*^*RNAi*^ (Fig. 4C, right traces). However, our data show that stj is required for normal pre-synaptic, somatodendritic, and axonal Ca_v_2 channel function, whereas dα_2_δ_1_ is not required for normal pre-synaptic Ca^2+^ channel function, but knock down increases functional axonal Ca_v_2 channel density.

**Figure 4:**
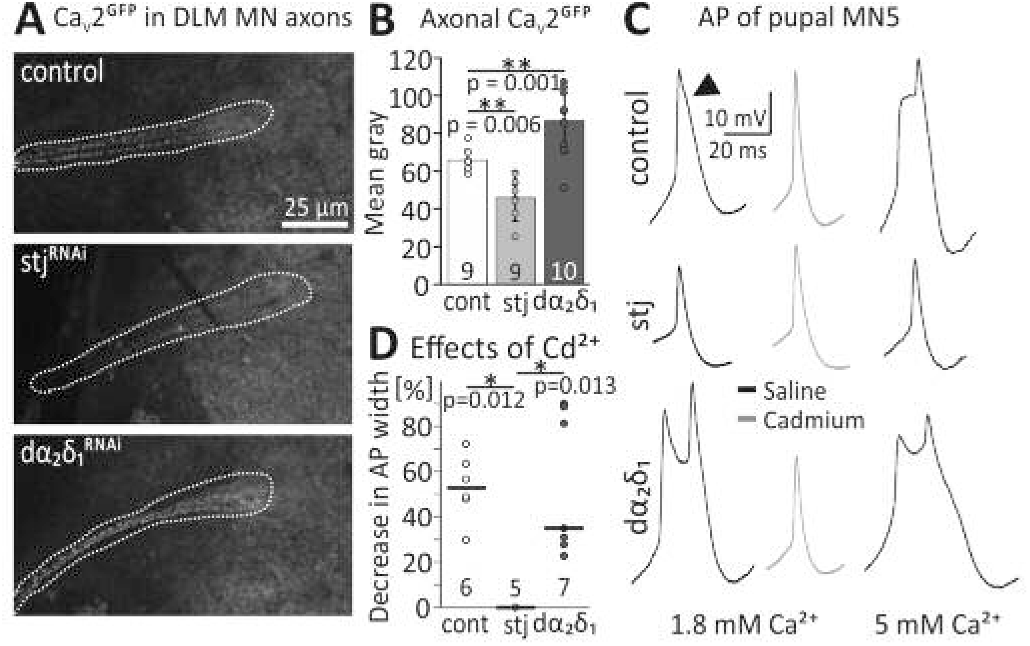
stj^RNAi^ and dα_2_δ_1_^RNAi^ affect normal function of axonal Ca_V_2 channels differently. **(A)** As compared to control (top panel) *stj*^*RNAi*^ (middle panel) decreases endogenous Ca_v_2^GFP^ label in DLM MN axons (encircled by dotted white line), whereas *dα*_*2*_*δ*_*1*_^*RNAi*^ (bottom panel) increases label. **(B)** Quantification of mean gray of Ca_v_2^GFP^ puncta in confocal sections revealed a ∼20% and significant decrease in *stj*^*RNAi*^ (light gray bar, **p=0.006, n=9) but a ∼20% and significant increase in *dα*_*2*_*δ*_*1*_^*RNAi*^ (dark gray bar, **p=0.001, n=10) as compared to control (white bar, N=9). Error bars represent SD; statistical significance was determined with one-way ANOVA with LSD post-hoc test. (**C**) Pupal DLM MN action potentials (APs) were recorded in 1.8 and 5 mM Ca^2+^ saline and elicited by square pulse current injections. APs showed a Ca^2+^ shoulder (C, top, left trace, see arrow) that was abolished by the VGCC blocker Cd^2+^ (500 µM; C, top, gray trace) and broadened in high Ca^2+^ (C, top, left trace). APs were smaller and narrower in *stj*^*RNAi*^ (**C**, middle, left two traces) as compared to control (**C**, top, left two traces) and were neither reduced or narrowed (**D**) nor broadened in high 5 mM Ca^2+^ saline (C, middle row, right trace). Following *dα*_*2*_*δ*_*1*_^*RNAi*^ APs often showed a double peak (**C**, bottom, left trace) that was abolished in Cd^2+^ (**C**, bottom, gray trace), but APs were broadened in high Ca^2+^ saline (**C**, bottom, left trace). AP half width was reduced significantly more in control (p = 0.012) and following *dα*_*2*_*δ*_*1*_^*RNAi*^ (p = 0.013) as compared to *stj*^*RNAi*^ (**D**, single data points and medians are presented. Statistical significance was determined by Kruskal-Wallis ANOVA with Dunn’s post-hoc test).

### stj and dα_2_δ_1_ are needed for dendritic localization of VGCCs, but only dα_2_δ_1_ sets axonal VGCC abundance

We next tested whether *stj*^*RNAi*^ and *dα*_*2*_*δ*_*1*_^*RNAi*^ affected also the functional localization of VGCCs in dendrites and axon. Somatodendritic Ca^2+^ currents were decreased following *stj*^*RNAi*^ but not following *dα*_*2*_*δ*_*1*_^*RNAi*^ (Fig. 3), and the Ca^2+^ component in pupal APs was abolished following *stj*^*RNAi*^ but increased following *dα*_*2*_*δ*_*1*_^*RNAi*^ (Fig. 4). Thus, we expected smaller dendritic and axonal Ca^2+^ influx in *stj*^*RNAi*^ MNs and unaltered dendritic but increased axonal Ca^2+^ influx following *dα_2_δ_1_^RNAi^*. We genetically expressed the Ca^2+^ indicator GCaMP6s (Chen et al., 2013, Ryglewski et al., 2017) in DLM MNs (23H06-GAL4>UAS-IVS-10xUAS-GCaMP6s) to assess potential channel mis-localization by functional imaging. We used pupal MNs because the developing VNC allows better visualization of dendritic processes as compared to the adult one, but overall localization of both dα_2_δ subunits through the VNC is similar during pupal and adult life. In these neurons, AP firing causes global Ca^2+^ influx through VGCC (Ryglewski et al., 2017) and was induced by somatic ramp current injection (Fig. 5C). The resulting Ca^2+^ signals were recorded from defined dendritic regions and from the axon (Fig. 5A). Unexpectedly, *stj* and *dα*_*2*_*δ*_*1*_ knock down both reduced dendritic Ca^2+^ signals by about 50% as compared to control (Figs. 5B, D) indicating a role of both dα_2_δ subunits for dendritic VGCC function. However, reduction in stj and dα_2_δ_1_ expression had opposing effects on axonal Ca^2+^ signals, as expected (Figs. 5B, Di). Axonal Ca^2+^ signals were significantly decreased following *stj*^*RNAi*^ but significantly increased following *dα_2_δ_1_^RNAi^*. Given that following *stj*^*RNAi*^ Ca^2+^ entry is reduced in all compartments tested, this indicates a principal role for stj to target and/or surface VGCCs or to increase channel conductance. However, a role of stj in increasing channel conductance seems unlikely as axonal Ca_v_2^GFP^ label is reduced in stj^RNAi^. By contrast, dα_2_δ_1_ specifically reduces dendritic but not axonal Ca^2+^ entry, which suggests a more specific role of dα_2_δ_1_ for targeting VGCCs to the dendritic domain. Reduced Ca_v_2 channel targeting to dendrites in *dα*_*2*_*δ*_*1*_^*RNAi*^ is further supported by concomitant dendritic growth defects (Fig. 6). It has previously been demonstrated that Ca_v_2 channels are required for local (Ryglewski et al., 2017) and for global dendritic growth regulation (Ryglewski et al., 2014; Vonhoff et al., 2013) in Drosophila DLM flight MNs. Similar to RNAi knock down of Ca_v_2 channels (Ryglewski et al., 2014) in the DLM MN5 *dα*_*2*_*δ*_*1*_^*RNAi*^ causes a significant decrease in total dendritic length (TDL, Fig. 6C) and in the number of branches (# branches, Fig. 6D) as revealed by intracellular dye fill and subsequent quantitative dendritic architecture analysis (Vonhoff and Duch, 2010). By contrast, mean dendritic branch length (MDL) and mean path length (MPL) are not affected (Figs. 6F-H). Therefore, *dα*_*2*_*δ*_*1*_^*RNAi*^ does not seem to affect dendritic territory borders or dendritic branch elongation, but causes a significant reduction in new dendritic branch formation or maintenance which results in reduced total length.

**Figure 5:**
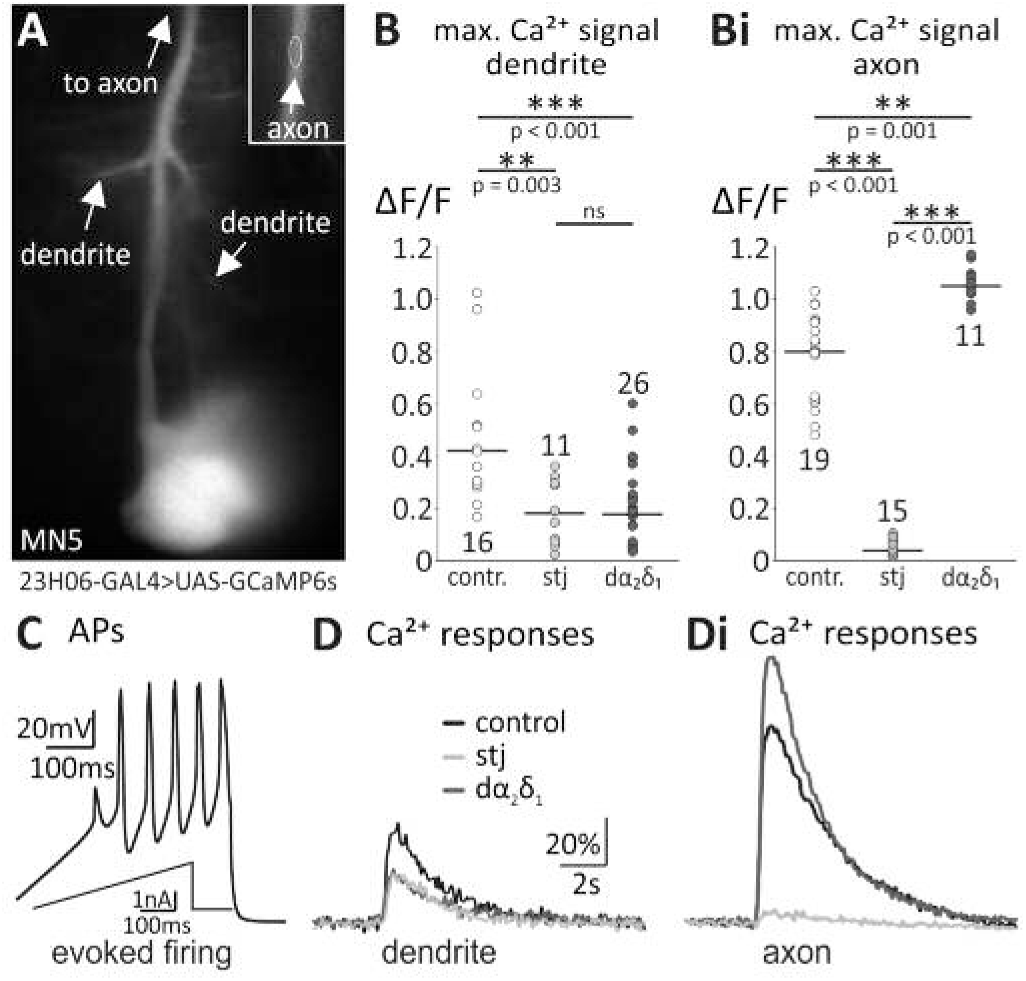
Dendritic localization of VGCCs depends on dα_2_δ_1_. (**A**) Changes in intracellular [Ca^2+^] (ΔF/F) upon induced firing in different compartments of pupal DLM MNs. Regions of interest in dendrites and in the axon (inset in **A**, upper right corner) are indicated by arrows and white circles. **(B, Bi)** Change in fluorescence in dendrites is significantly reduced following both *stj*^*RNAi*^ (light gray circles; **p=0.003, n=11) and *dα*_*2*_*δ*_*1*_^*RNAi*^ (dark gray circles; ***p<0.001, n=26) as compared to control (**B**, white circles, n=16), whereas no statistical difference is found between *stj*^*RNAi*^ and *dα*_*2*_*δ*_*1*_^*RNAi*^ (ns; **B**). By contrast, axonal Ca^2+^ signals are significantly reduced following *stj*^*RNAi*^ (light gray circles; ***p<0.001, n=15) but significantly increased following *dα*_*2*_*δ*_*1*_^*RNAi*^ (**p=0.001, N=11) as compared to control (**Bi**, N=19). Change in fluorescence is significantly reduced following *stj*^*RNAi*^ as compared to *dα*_*2*_*δ*_*1*_^*RNAi*^ (***p<0.001; **Bi**). Data in (**B** and **Bi)** are presented as single data points and median. Statistical significance was tested by Kruskal-Wallis ANOVA with Dunn’s post-hoc test. (**C-Di**) Change in fluorescence (ΔF/F) upon evoked AP firing in different neuronal compartments in control and following *stj*^*RNAi*^ and *dα_2_δ_1_^RNAi^*. (**C**) APs were elicited by somatic ramp current injection of 400 ms duration and 1 nA amplitude (stimulation protocol see inset in **C**). (**D**) Dendritic Ca^2+^ signals are reduced approx. by half following both *stj*^*RNAi*^ (light gray trace) and *dα*_*2*_*δ*_*1*_^*RNAi*^ (dark gray trace) as compared to control (black trace). (**Di**) Axonal Ca^2+^ signals are reduced following *stj*^*RNAi*^ (light gray trace) but increased following *dα*_*2*_*δ*_*1*_^*RNAi*^ (light gray trace) as compared to control (black trace).

**Figure 6:**
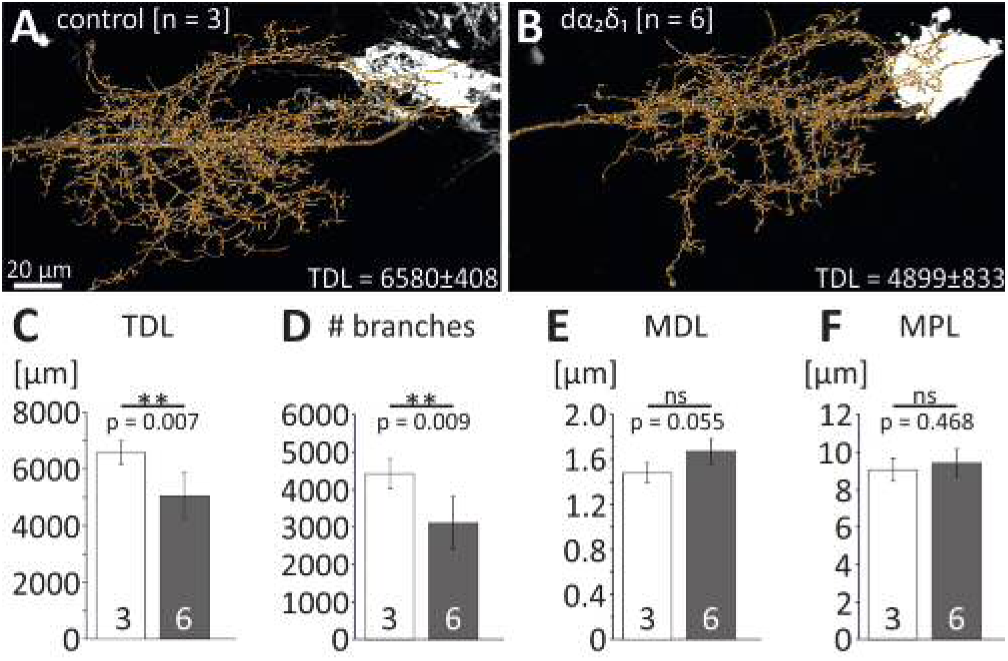
dα_2_δ_1_ affects dendrite development. **(A, B)** Reconstructions of adult MN5 dendrite in control (**A**, n=3) and following *dα*_*2*_*δ*_*1*_^*RNAi*^ (**B**, n=6). **(C-F)** Morphometric parameters were analyzed. Following *dα*_*2*_*δ*_*1*_^*RNAi*^ total dendritic length (**C**, TDL, 6580 ± 408 µm vs. 5043 ± 824 µm; **p = 0.007, Student’s T-test) as well as the number of dendritic branches (**D**, # branches, 4421 ± 382 vs. 3122 ± 700; **p=0.009, Student’s T-test) are reduced. Other parameters like the mean dendrite length (**E**, MDL, 1.48 ± 0.09 vs. 1.67 ± 0.11 µm; p=0.055) and the mean path length (**F**, MPL, 9.05 ± 0.61 vs. 9.41 ± 0.74 µm; p=0.468) are not affected.

## Discussion

In this study we find that dα_2_δ_1_ and dα_2_δ_3_ (stj) are broadly expressed in the Drosophila nervous system. Despite differential expression patterns through the ventral nerve cord, both are co-expressed in the same motoneurons where they localize to different sub-neuronal compartments and mediate different functions. dα_2_δ_3_ (stj) is required for normal motoneuron presynaptic function and full somatodendritic and axonal Ca_v_1 and Ca_v_2 current amplitudes. By contrast, dα_2_δ_1_ is not required for pre-synaptic function or full calcium current amplitudes, but by contrast, for the correct relative Ca_v_2 channel allocation to the dendritic versus the axonal domain. Therefore, our *in vivo* analysis indicates that different α_2_δ subunits are required in the same neurons to regulate distinctly different functions of voltage gated Ca^2+^ channels. This contrasts findings in heterologous expression systems, where any HVA channel requires co-expression of any α_2_δ subunit for proper current amplitudes (Dolphin, 2012), but it is in agreement with different brain diseases resulting from specific mutations of single α_2_δ subunits. This underscores the importance for *in vivo* studies to unravel the combinatorial code by which different α_2_δ/α_1_ interactions mediate functional Ca^2+^ channel diversity in different types of neurons and different sub-neuronal compartments.

### dα_2_δ_1_ and dα_2_δ_3_ (stj) mediate distinctly different functions in the CNS and in the same neurons

Mutations in single mammalian *α*_*2*_*δ* genes are known causes to multiple brain diseases (Calandre et al., 2016; Davies et al., 2007; Klugbauer et al., 2003), thus indicating that loss of function of one α_2_δ subunit cannot be compensated for by others. Similarly, *dα*_*2*_*δ*_*3*_ null mutant animals are embryonic lethal (Dickman et al., 2008; Ly et al., 2008; Kurshan et al., 2009). In addition, we find that *dα*_*2*_*δ*_*3*_^*RNAi*^ targeted specifically to flight motoneurons causes inability to fly, *dα*_*2*_*δ*_*1*_^*RNAi*^ significantly reduces Drosophila climbing speed, and pan-neural RNAi knock down of either or *dα*_*2*_*δ*_*3*_ or *dα*_*2*_*δ*_*1*_ does not cause compensatory up-regulation of the expression levels of the other one. Together these findings indicate differential and non-redundant functions of *dα*_*2*_*δ*_*1*_ and *dα*_*2*_*δ*_*3*_ (*stj*). On the contrary, in heterologous expression systems co-expression of multiple different α_2_δ subunits each can support the same HVA current (Dolphin, 2012), indicating that in principle, different α_1_ / α_2_δ combinations can mediate similar functions, at least with regard to channel biophysical properties and surfacing. However, *in vivo*, redundant functions of multiple different α_1_ / α_2_δ combinations seems unlikely, because differential spatial expression of different α_2_δ subunits has been reported for vertebrate brain (Scott and Kammermeier 2017; Nieto-Rostro et al., 2014; Cole et al., 2005). Similarly, we find differential expression patterns of dα_2_δ_1_ and stj in the Drosophila ventral nerve cord. dα_2_δ_1_ is strongly expressed in all central neuropils and in many neuronal somata, whereas dα_2_δ_3_ was not detected in central neuropils but in many neuronal somata and in motoneuron axon terminals at the neuromuscular junction (Dickman et al., 2008; Ly et al., 2008; Kurshan et al., 2009). Notably, in Drosophila motoneurons both dα_2_δ_1_ and dα_2_δ_3_ are co-expressed, but localize to different sub-neuronal compartments. dα_2_δ_3_ but not dα_2_δ_1_ localizes at axon terminals and is required for pre-synaptic function (Wang et al., 2016; this study) and embryonic synapse development (Kurshan et al., 2009). Moreover, we find that dα_2_δ_3_ and dα_2_δ_1_ affect different aspects of motoneuron calcium channel function (see below).

### dα_2_δ_3_ (stj) is required for normal Ca_v_1 and Ca_v_2 currents in all sub-neuronal compartments

*dα*_*2*_*δ*_3_ has previously been reported essential for Drosophila neuromuscular synapse development and function. During embryonic development loss of dα_2_δ_3_ (stj) function impairs early steps of synapse formation before calcium channels arrive at the presynaptic terminal, as well as the subsequent recruitment of Ca_v_2-like channels to the active zone (Kurshan et al., 2009). Moreover, at mature larval neuromuscular junctions, dα_2_δ_3_ is required for rapid induction and continuous expression of pre-synaptic homeostatic potentiation (Wang et al., 2016). Our data demonstrate that dα_2_δ_3_ is not only required for Ca_v_2 channel function at the pre-synaptic terminal, but also for normal flight motoneuron somatodendritic and axonal calcium currents, both of which are mediated by the Drosophila Ca_v_2 channel homolog cacophony (Ryglewski et al., 2012; 2014). In addition, dα_2_δ_3_ is required for normal larval motoneuron somatodendritic Ca_v_1 current amplitudes encoded by the L-type channel homolog *Dmca1D* (Worrell and Levine, 2008; Kadas et al., 2017; Schützler et al., 2019). Therefore, dα_2_δ_3_ (stj) is required for normal calcium current amplitudes in Drosophila motoneurons independent of sub-neuronal compartment or Ca_v_1 or Ca_v_2 channel, thus indicating a general function in Ca_v_1 and Ca_v_2 channel surfacing. An important role for channel surfacing has also been reported for vertebrate α_2_δ_1_ (D’Arco et al., 2015; Cassidy et al., 2014).

### α_2_δ_1_ specifically sets dendritic calcium channel abundance

Although somatodendritic calcium current amplitude is reduced by *dα*_*2*_*δ*_*3*_^*RNAi*^ but unaffected by *dα*_*2*_*δ*_*1*_^*RNAi*^, our data indicate that dα_2_δ_1_ affects the relative allocation of calcium channels to the axonal versus the dendritic domain. Please note that voltage clamp recordings from the soma reflect the sum of proximal and more distal currents, but cannot precisely detect relative shifts in channel localization. Functional imaging reveals that both, targeted RNAi knock down of *dα*_*2*_*δ*_*1*_ and *dα*_*2*_*δ*_*3*_ significantly reduce the amplitude of dendritic calcium signals as evoked by action potential firing. By contrast, axonal calcium signals as evoked by the same firing patterns are nearly eliminated by *dα*_*2*_*δ*_*3*_^*RNAi*^ but significantly increased by *dα_2_δ_1_*^RNAi^. This again demonstrates different functions of *dα*_2_δ_1_ and *dα*_*2*_*δ*_*3*_ in the same neurons. Reduced dendritic and axonal calcium signals upon *dα*_*2*_*δ*_*3*_^*RNAi*^ down are in line with a general function of stj in channel surfacing (see above). By contrast, increased axonal but decreased dendritic calcium signals following *dα*_*2*_*δ*_*1*_^*RNAi*^ are in accord with a function of dα_2_δ_1_ in channel sorting and/or targeting. One possibility is that dα_2_δ_1_ serves linkage to specific motor proteins for transport to the dendritic domain. To us, the most parsimonious explanation for increased axonal calcium current amplitudes is that at a given rate of Ca_v_2 channel production, reduced transport to dendrites in *dα*_*2*_*δ*_*1*_^*RNAi*^ leaves increased channel numbers for the axon. By contrast, in mouse sensory neurons, knock-out of *α*_*2*_*δ*_*1*_ decreases axonal calcium channels (Margas et al., 2016). Please note that dα_2_δ_1_ does not necessarily correspond to vertebrate α_2_δ_1_. Although Drosophila α_2_δ_1_ and α_2_δ_3_ contain the essential functional domains of vertebrate α_2_δ subunits, like the MIDAS motif, the von Willebrandt factor A (VWA), and the cache domains, sequence homology is not high enough to unambiguously match each vertebrate α_2_δ subunit with a specific Drosophila one. Based on functional analysis so far available one might speculate that vertebrate α_2_δ_1_ and Drosophila dα_2_δ_3_ are functional pendants, because both are required for Ca_v_2 channel targeting to axon terminals (Kurshan et al., 2009; Hoppa et al., 2012), both increase calcium channel abundance in the axonal membrane (this study; Margas et al., 2016), both increase Cav2 current amplitude (this study; D’Arco et al., 2015), and both play roles in the development of excitatory synapses independent of calcium channel function (Kurshan et al., 2009; Eroglu et al., 2009; Risher et al., 2018).

## Acknowledgments

Support from the German Research Foundation (Deutsche Forschungsgemeinschaft, DFG) to SR is gratefully acknowledged (RY117/3-1). We thank Dr. Duch (Mainz) for many helpful discussions and comments on the ms.

The authors declare no conflict of interest.

---

**Table S1.** Fly stocks used in this study

| Genotype | use | identifier |
| --- | --- | --- |
| y[1] w[67c23]; Mi{PT-GFSTF.0}CG4587[Mi01722-GFSTF.0] | $\alpha_2\delta_1$ protein trap, endogenously expressing GFP fused to $\alpha_2$ fragment of $\alpha_2\delta_1$, resulting in $\alpha_2\delta_1^{GFP}$ (Nagarkar-Jaiswal et al., 2015) | RRID:BDSC_59288 |
| y[1] w[*]; Mi{PT-GFSTF.0}stj[Mi00783-mCherry.0]/CyO Tb | stj protein trap, endogenously expressing mCherry fused to $\alpha_2$ fragment of stj, resulting in $stj^{mCherry}$ | this study |
| w[1118]; P{KK106795}VIE-260B; P{y[+t7.7] v[+t1.8]=TRiP.JF01825}attP2 | UAS-$\alpha_2\delta_1^{RNAiKK}$ with UAS-stj<sup>RNAi</sup> in the VALIUM10 vector (Perkins et al., 2015) | RRID:FlyBase_FBst0479962;<br>RRID:BDSC_25807 |
| y* w*; P{GawB}stjNP1574 / CyO, P{UAS-lacZ.UW14}UW14 | stj-GAL4 (Ly et al., 2008) | RRID:DGGR_104034 |
| w[1118]; P{KK101267}VIE-260B; P{w[+mC]=UAS-Dcr-2.D}10 | UAS-stj<sup>RNAiKK</sup> with UAS-dcr2 to enhance RNAi efficacy (Dietzl et al., 2007) | RRID:FlyBase_FBst0480379;<br>RRID:BDSC_24651 |
| w[1118]; P{KK106795}VIE-260B; P{w[+mC]=UAS-Dcr-2.D}10 | UAS-$\alpha_2\delta_1^{RNAiKK}$ with UAS-dcr2 to enhance RNAi efficacy (Dietzl et al., 2007) | RRID:FlyBase_FBst0479962;<br>RRID:BDSC_24651 |
| w[1118]; P{w[+mW.hs]=GawB}VGlut[OK371] | vGlut-GAL4, expressed in all motoneurons incl. larval crawling MNs | RRID:BDSC_26160 |
| w[1118]; P{y[+t7.7] w[+mC]=20XUAS-IVS-GCaMP6s}attP40; P{y[+t7.7] w[+mC]=GMR23H06-GAL4}attP2 | UAS-GCaMP6s is a genetically encodes $Ca^{2+}$ indicator. 23H06-GAL4 from HHMI Janelia Farm GAL4 driver lines; associated gene CG8084 *anachronism*, for expression pattern see http://flweb.janelia.org/cgi-bin/flew.cgi (R23H06). 23H06-Gal4 expresses in pupal and adult DLM MNs and only very few others | RRID:BDSC_42746;<br>23H06-GAL4 discontinued at BDSC (available from our lab) |
| P{w[+mW.hs]=GawB}elav[C155] | pan-neural GAL4-driver, used for pan-neural expression of UAS-RNAi transgenes for Western Blot analysis and climbing assay | RRID:BDSC_458 |
| w[1118]; P{w[+mC]=UAS-Dcr-2.D}2; P{y[+t7.7] w[+mC]=GMR23H06-GAL4}attP2<br>P{w[+mC]=UAS-myr-mRFP}2/TM6B, Tb[1] | UAS-dcr2 to enhance RNAi efficacy (Dietzl et al., 2007) along with 23H06-GAL4 with UAS-myr-mRFP on the same chromosome (Chr. 3) for easier identification of DLM MNs. for expression. Used for expression of double RNAi of stj and $\alpha_2\delta_1$ | RRID:BDSC_24650;<br>23H06-GAL4 discontinued at BDSC (available from our lab) |
| w[1118]; P{y[+t7.7] w[+mC]=GMR23H06- | 23H06-GAL4 along with UAS-myr- | 23H06-GAL4 discontinued at BDSC |
| GAL4}attP2, P{w[+mC]=UAS-myr-mRFP}2/TM6B, Tb[1] | mRFP on the same chromosome (Chr. 3) for easier identification of DLM MNs. | (available from our lab);<br>RRID:BDSC_7119 |
| P{w[+mW.hs]=GawB}elav[C155], *cac<sup>sfGFP-N</sup>* | GFP tagged cacophony along with *elav<sup>C155</sup>*-GAL4 on the same chromosome. cacophony VGCCs are endogenously tagged with super folder (sf) GFP. GFP tag was inserted directly after the second translational start via CRISPR/cas9-based methods (Gratz et al., 2019). | RRID:BDSC_458;<br>identifier for *cac<sup>sfGFP-N</sup>* not yet assigned (Gratz et al. 2019) |
| w[*]; P{w[+mC]=eve-GAL4.RN2}P,<br>P{w[+mC]=UAS-mCD8::GFP.L}LL5/CyO;<br>P{w[+mC]=Act(FRT.stop)GAL4},<br>P{w[+mC]=UAS-FLP.D}JD2 | inducible FLP out in larval MN1s and MN1b crawling MNs to induce mosaic expression of UAS.-transgenes after second instar (eve-GAL4 is turned off before third instar) | RRID:BDSC_7475;<br>RRID:BDSC_4540;<br>fly strain courtesy of Dr. S. Sanyal, Calico labs, San Francisco |
| w[*];P{w[+mC]=UAS-mCD8::GFP.L}LL5;<br>P{w[+mW.hs]=GawB}D42 | GAL4-expression in motoneurons (incl. DLM MNs) under the control of the toll6 receptor | RRID:BDSC_8816 |
| P{w[+mC]=UAS-FLAG-stj-HA}2/CyO; In(3L)D,<br>D[1]/TM3, Sb[1] | HA-tagged transgene of stj | RRID:BDSC_39717 |
| y[1] M{vas-int.B}ZH-2A w[*]; sna[Sco]/CyO,<br>P{ry[+t7.2]=sevRas1.V12}FK1 | φC31 integrase expression under the control of *vasa*, which is active in the germ line | RRID:BDSC_36312 |
| y[1] w[*];<br>Mi{y[+mDint2]=MIC}stj[MI00783]/SM6a | fly strain carrying a MiMIC within a stj intron | RRID:BDSC_34109 |
| y[1] w[*];CyO/sna[Sco] | balancer strain | lab stock |

**Table S2.**
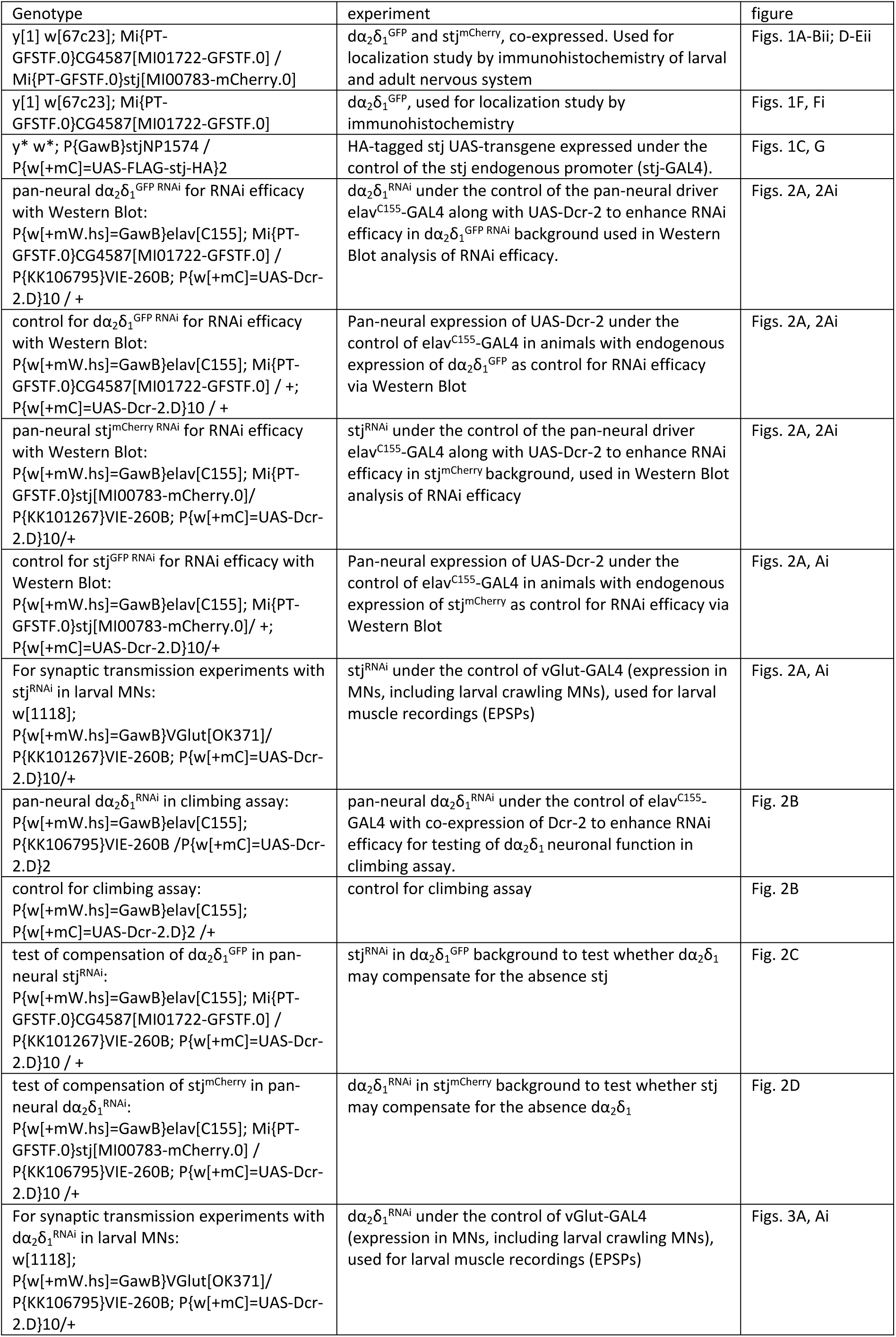
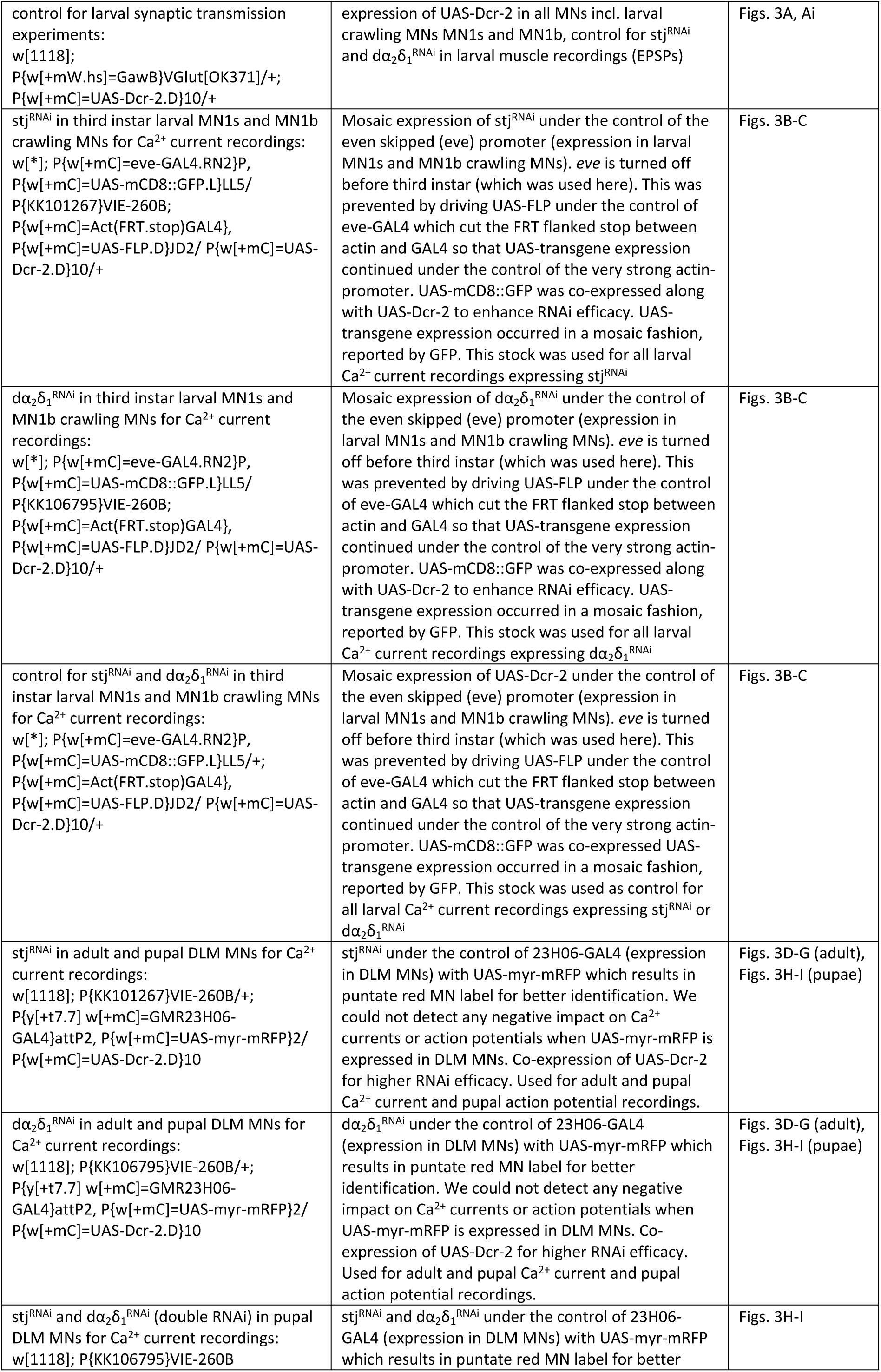
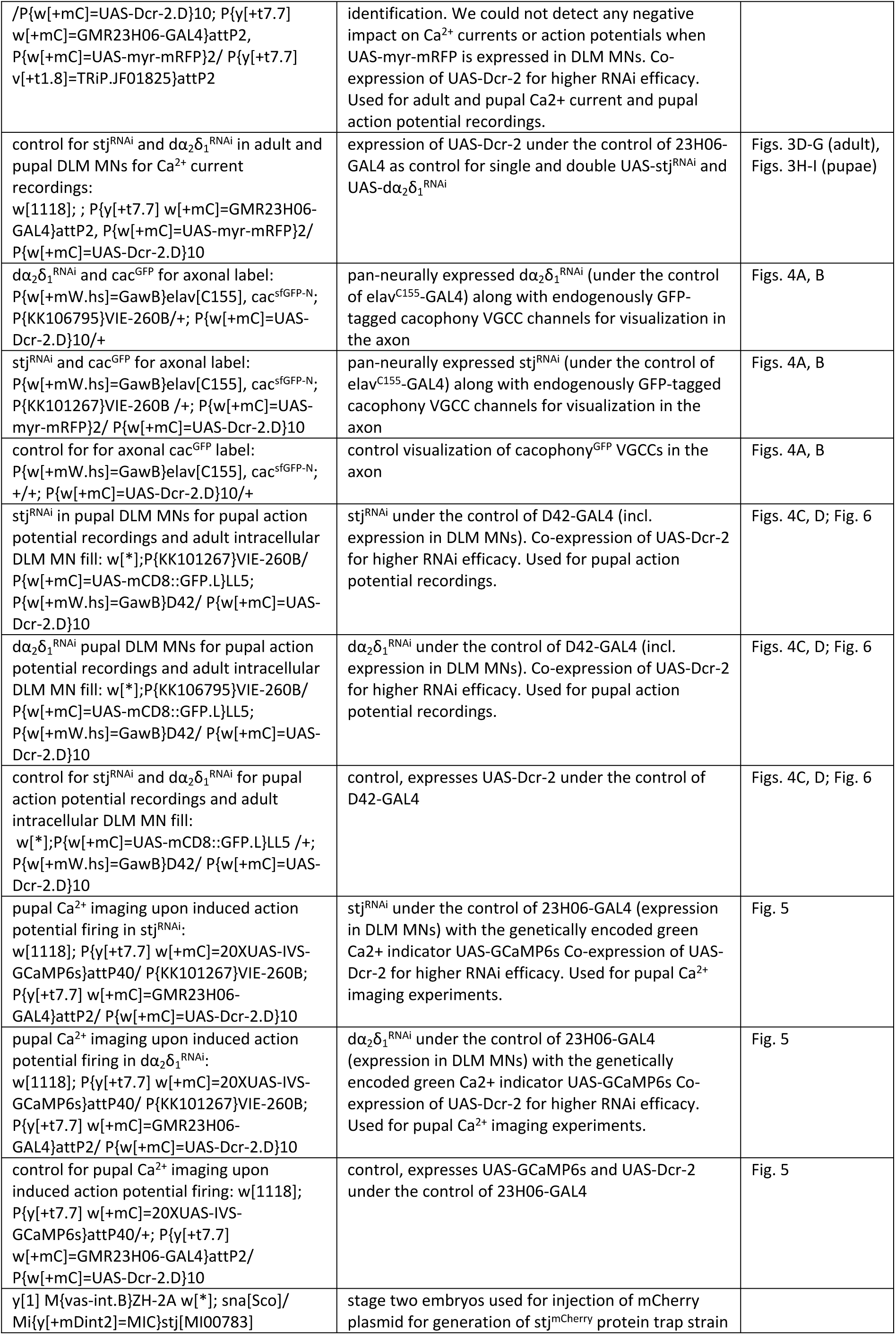
Genotypes used for experiments

## References

Bainbridge SP, Bownes M (1981) Staging the metamorphosis of Drosophila melanogaster. J Embryol Exp Morphol 66:57–80.

Barclay J, Balaguero N, Mione M, Ackerman SL, Letts VA, Brodbeck J, Canti C, Meir A, Page KM, Kusumi K, Perez-Reyes E, Lander ES, Frankel WN, Gardiner RM, Dolphin AC, Rees M (2001) Ducky mouse phenotype of epilepsy and ataxia is associated with mutations in the Cacna2d2 gene and decreased calcium channel current in cerebellar Purkinje cells. J Neurosci 21(16):6095–6104.

Bauer CS, Nieto-Rostro M, Rahman W, Tran-Van-Minh A, Ferron L, Douglas L, Kadurin I, Sri Ranjan Y, Fernandez-Alacid L, Millar NS, Dickenson AH, Lujan R, Dolphin AC (2009) The increased trafficking of the calcium channel subunit alpha2delta-1 to presynaptic terminals in neuropathic pain is inhibited by the alpha2delta ligand pregabalin. J Neurosci 29(13):4076–4088.

Brockhaus J, Schreitmüller M, Repetto D, Klatt O, Reissner C, Elmslie K, Heine M, Missler M (2018) α-Neurexins Together with α2δ-1 Auxiliary Subunits Regulate Ca2+ Influx through Cav2.1 Channels. J Neurosci. 38(38):8277–8294.

Brodbeck J, Davies A, Courtney JM, Meir A, Balaguero N, Canti C, Moss FJ, Page KM, Pratt WS, Hunt SP, Barclay J, Rees M, Dolphin AC (2002), The ducky mutation in Cacna2d2 results in altered Purkinje cell morphology and is associated with the expression of a truncated alpha 2 delta-2 protein with abnormal function. J Biol Chem 277(10):7684–7693.

Buraei Z, Yang J (2010), The ß subunit of voltage-gated Ca2+ channels. Physiol Rev. 90(4):1461–1506. Review.

Cantí C, Nieto-Rostro M, Foucault I, Heblich F, Wratten J, Richards MW, Hendrich J, Douglas L, Page KM, Davies A, Dolphin AC (2005) The metal-ion-dependent adhesion site in the Von Willebrand factor-A domain of alpha2delta subunits is key to trafficking voltage-gated Ca2+ channels. Proc. Natl. Acad. Sci. U.S.A. 102(32):11230–11235.

Calandre EP, Rico-Villademoros F, Slim M (2016) Alpha2delta ligands, gabapentin, pregabalin and mirogabalin: a review of their clinical pharmacology and therapeutic use. Expert Rev Neurother. 16(11):1263–1277. Review. Erratum in Expert Rev Neurother. 16(11):iii (2016).

Cassidy JS, Ferron L, Kadurin I, Pratt WS, Dolphin AC (2014) Functional exofacially tagged N-type calcium channels elucidate the interaction with auxiliary α2δ-1 subunits. Proc Natl Acad Sci U.S.A. 111(24):8979–8984.

Catterall WA (2011) Voltage-gated calcium channels. Cold Spring Harb Perspect Biol. 3(8):a003947. doi: 10.1101/cshperspect.a003947. Review.

Celli R, Santolini I, Guiducci M, van Luijtelaar G, Parisi P, Striano P, Gradini R, Battaglia G, Ngomba RT, Nicoletti F (2017) The α2δ Subunit and Absence Epilepsy: Beyond Calcium Channels? Curr Neuropharmacol. 15(6):918–925. Review.

Chen Y, Chen SR, Chen H, Zhang J, Pan HL (2019) Increased α2δ-1-NMDA receptor coupling potentiates glutamatergic input to spinal dorsal horn neurons in chemotherapy-induced neuropathic pain. J Neurochem. 148(2):252–274.

Chen TW, Wardill TJ, Sun Y, Pulver SR, Renninger SL, Baohan A, Schreiter ER, Kerr RA, Orger MB, Jayaraman V, Looger LL, Svoboda K, Kim DS (2013) Ultrasensitive fluorescent proteins for imaging neuronal activity. Nature. 499(7458):295–300.

Cole RL, Lechner SM, Williams ME, Prodanovich P, Bleicher L, Varney MA, Gu G (2005) Differential Distribution of Voltage-Gated Calcium Channel Alpha-2 Delta (α2δ) Subunit mRNA-Containing Cells in the Rat Central Nervous System and the Dorsal Root Ganglia. J Comp. Neurol. 491:246–269.

D’Arco M, Margas W, Cassidy JS, Dolphin AC (2015) The upregulation of α2δ-1 subunit modulates activity-dependent Ca2+ signals in sensory neurons. J Neurosci. 35(15):5891–5903.

Davies A, Douglas L, Hendrich J, Wratten J, Tran-Van-Minh A, Foucault I, Koch D, Pratt WS, Saibil H, Dolphin AC (2006) The calcium channel α2δ-2 subunit partitions with CaV2.1 in lipid rafts in cerebellum: implications for localization and function, J Neurosci. 26:8748–8757.

Davies A, Hendrich J, Van Minh AT, Wratten J, Douglas L, Dolphin AC (2007) Functional biology of the alpha(2)delta subunits of voltage-gated calcium channels. Trends Pharmacol Sci. 28(5):220–228. Review.

Davies A, Kadurin I, Alvarez-Laviada A, Douglas L, Nieto-Rostro M, Bauer CS, Pratt WS, Dolphin AC (2010) The α2δ subunits of voltage-gated calcium channels form GPI anchored proteins, a posttranslational modification essential for function. Proc. Natl. Acad. Sci. U.S.A. 107(4):1654–1659.

Dickman DK, Kurshan PT, Schwarz TL (2008) Mutations in a Drosophila alpha2delta voltage-gated calcium channel subunit reveal a crucial synaptic function. J Neurosci. 28(1):31–38.

Dietzl G, Chen D, Schnorrer F, Su KC, Barinova Y, Fellner M, Gasser B, Kinsey K, Oppel S, Scheiblauer S, Couto A, Marra V, Keleman K, Dickson BJ (2007) A genome-wide transgenic RNAi library for conditional gene inactivation in Drosophila. Nature. 448(7150):151–156.

Dolphin AC (2012) Calcium channel α2δ subunits in epilepsy and as targets for antiepileptic drugs” in Jasper’s Basic Mechanisms of the Epilepsies [Internet]. J. L. Noebels, M. Avoli, M. A. Rogawski, R. W. Olsen, A. V. Delgado-Escueta, Eds. 4th edition. (Bethesda (MD): National Center for Biotechnology Information (US), 2012).

Dolphin AC (2013) The α2δ subunits of voltage-gated calcium channels. Biochim Biophys Acta. 1828(7):1541–1549.

Eroglu C, Allen NJ, Susman MW, O’Rourke NA, Park CY, Ozkan E, Chakraborty C, Mulinyawe SB, Annis DS, Huberman AD, Green EM, Lawler J, Dolmetsch R, Garcia KC, Smith SJ, Luo ZD, Rosenthal A, Mosher DF, Barres BA (2009) Gabapentin receptor alpha2delta-1 is a neuronal thrombospondin receptor responsible for excitatory CNS synaptogenesis. Cell 139(2):380–392.

Evers JF, Schmitt S, Sibila M, Duch C (2005) Progress in functional neuroanatomy: precise automatic geometric reconstruction of neuronal morphology from confocal image stacks. J Neurophysiol 93(4):2331–2342.

Faria LC, Gu F, Parada I, Barres B, Luo ZD, Prince DA (2017) Epileptiform activity and behavioral arrests in mice overexpressing the calcium channel subunit α2δ-1. Neurobiol Dis. 102:70–80.

Felix R, Gurnett CA, De Waard M, Campbell KP (1997) Dissection of functional domains of the voltage-dependent Ca2+ channel alpha2delta subunit. J Neurosci 17(18):6884–6891.

Fell B, Eckrich S, Blum K, Eckrich T, Hecker D, Obermair GJ, Münkner S, Flockerzi V, Schick B, Engel J (2016) α2δ2 Controls the Function and Trans-Synaptic Coupling of Cav1.3 Channels in Mouse Inner Hair Cells and Is Essential for Normal Hearing. J Neurosci. 36(43):11024–11036.

Feng Y, Ueda A, Wu CF (2004) A modified minimal hemolymph-like solution, HL3.1, for physiological recordings at the neuromuscular junctions of normal and mutant Drosophila larvae. J Neurogenet 18(2):377–402.

Ferron L, Kadurin I, Dolphin AC (2018) Proteolytic maturation of α2δ controls the probability of synaptic vesicular release. Elife 7. pii: e37507 (2018). doi: 10.7554/eLife.37507.

Gratz SJ, Goel P, Bruckner JJ, Hernandez RX, Khateeb K, Macleod GT, Dickman D, O’Connor-Giles KM (2019) Endogenous Tagging Reveals Differential Regulation of Ca2+ Channels at Single Active Zones during Presynaptic Homeostatic Potentiation and Depression. J Neurosci 39(13):2416–2429.

Green EW, Fedele G, Giorgini F, Kyriacou CP (2014) A Drosophila RNAi collection is subject to dominant phenotypic effects. Nat Methods 11(3):222–223.

Hendrich J, Tran-Van-Minh A, Heblich F, Nieto-Rostro M, Watschinger K, Striessnig J, Wratten J, Davies A, Dolphin AC (2008) Pharmacological disruption of calcium channel trafficking by the α2δ ligand gabapentin, Proc. Natl. Acad. Sci. U.S.A. 105:3628–3633.

Hobom M, Dai S, Marais E, Lacinova L, Hofmann F, Klugbauer N (2000) Neuronal distribution and functional characterization of the calcium channel alpha2delta-2 subunit. Eur J Neurosci 12(4):1217–1226.

Hoppa MB, Lana B, Margas W, Dolphin AC, Ryan TA (2012) α2δ expression sets presynaptic calcium channel abundance and release probability. Nature 486(7401):122–125.

Kadas D, Klein A, Krick N, Worrell JW, Ryglewski S, Duch C (2017) Dendritic and Axonal L-Type Calcium Channels Cooperate to Enhance Motoneuron Firing Output during Drosophila Larval Locomotion. J Neurosci 37(45):10971–10982.

Kadurin I, Rothwell SW, Lana B, Nieto-Rostro M, Dolphin AC (2017) LRP1 influences trafficking of N-type calcium channels via interaction with the auxiliary α2δ-1 subunit. Sci Rep 7:43802 (2017). doi: 10.1038/srep43802.

Kawasaki F, Zou B, Xu X, Ordway RW (2004) Active zone localization of presynaptic calcium channels encoded by the cacophony locus of Drosophila. J Neurosci 24(1):282–285.

Kittel RJ, Wichmann C, Rasse TM, Fouquet W, Schmidt M, Schmid A, Wagh DA, Pawlu C, Kellner RR, Willig KI, Hell SW, Buchner E, Heckmann M, Sigrist SJ, Bruchpilot promotes active zone assembly, Ca2+ channel clustering, and vesicle release. Science 312(5776):1051–1054.

Klugbauer N, Marais E, Hofmann F (2003) Calcium channel alpha2delta subunits: differential expression, function, and drug binding. J Bioenerg Biomembr 35(6):639–647. Review.

Kurshan PT, Oztan A, Schwarz TL (2009) Presynaptic alpha2delta-3 is required for synaptic morphogenesis independent of its Ca2+-channel functions. Nat Neurosci 12(11):1415–1423.

Littleton JT, Ganetzky B (2000) Ion channels and synaptic organization: analysis of the Drosophila genome. Neuron 26(1):35–43.

Ly CV, Yao CK, Verstreken P, Ohyama T, Bellen HJ (2008) straightjacket is required for the synaptic stabilization of cacophony, a voltage-gated calcium channel alpha1 subunit. J Cell Biol 181(1):157–70.

Margas W, Ferron L, Nieto-Rostro M, Schwartz A, Dolphin AC (2016) Effect of knockout of α2δ-1 on action potentials in mouse sensory neurons. Philos Trans R Soc Lond B Biol Sci. 371(1700) (2016). pii: 20150430. doi: 10.1098/rstb.2015.0430.

Nagarkar-Jaiswal S, Lee PT, Campbell ME, Chen K, Anguiano-Zarate S, Cantu Gutierrez M, Busby T, Lin WW, He Y, Schulze KL, Booth BW, Evans-Holm M, Venken KJ, Levis RW, Spradling AC, Hoskins RA, Bellen HJ (2015) A library of MiMICs allows tagging of genes and reversible, spatial and temporal knockdown of proteins in Drosophila. eLife 4: e05338 (2015). doi:10.7554/eLife.05338.

Nieto-Rostro M, Ramgoolam K, Pratt WS, Kulik A, Dolphin AC (2018) Ablation of α2δ-1 inhibits cell-surface trafficking of endogenous N-type calcium channels in the pain pathway in vivo. Proc Natl Acad Sci U.S.A. 115(51):E12043–E12052.

Nieto-Rostro M, Sandhu G, Bauer CS, Jiruska P, Jefferys JG, Dolphin AC (2014) Altered expression of the voltage-gated calcium channel subunit α₂δ-1: a comparison between two experimental models of epilepsy and a sensory nerve ligation model of neuropathic pain. Neuroscience 283:124–137.

Perkins LA, Holderbaum L, Tao R, Hu Y, Sopko R, McCall K, Yang-Zhou D, Flockhart I, Binari R, Shim HS, Miller A, Housden A, Foos M, Randkelv S, Kelley C, Namgyal P, Villalta C, Liu LP, Jiang X, Huan-Huan Q, Wang X, Fujiyama A, Toyoda A, Ayers K, Blum A, Czech B, Neumuller R, Yan D, Cavallaro A, Hibbard K, Hall D, Cooley L, Hannon GJ, Lehmann R, Parks A, Mohr SE, Ueda R, Kondo S, Ni JQ, Perrimon N (2015) The Transgenic RNAi Project at Harvard Medical School: Resources and Validation. Genetics 201(3): 843–852.

Reuveny A, Shnayder M, Lorber D, Wang S, Volk T (2018) Ma2/d promotes myonuclear positioning and association with the sarcoplasmic reticulum. Development 28:145(17). doi: 10.1242/dev.159558.

Risher WC, Kim N, Koh S, Choi JE, Mitev P, Spence EF, Pilaz LJ, Wang D, Feng G, Silver DL, Soderling SH, Yin HH, Eroglu C (2018) Thrombospondin receptor α2δ-1 promotes synaptogenesis and spinogenesis via postsynaptic Rac1. J Cell Biol 217(10):3747–3765.

Ryglewski S, Duch C (2012b) Preparation of Drosophila central neurons for in situ patch clamping. J Vis Exp. 2012 Oct 15;(68). pii: 4264. doi: 10.3791/4264.S.

Ryglewski S, Kilo L, Duch C (2014) Sequential acquisition of cacophony calcium currents, sodium channels and voltage-dependent potassium currents affects spike shape and dendrite growth during postembryonic maturation of an identified Drosophila motoneuron. Eur J Neurosci 39(10):1572–1585.

Ryglewski S, Lance K, Levine RB, Duch C (2012) Ca(v)2 channels mediate low and high voltage-activated calcium currents in Drosophila motoneurons. J Physiol 590(4):809–258.

Ryglewski S, Vonhoff F, Scheckel K, Duch C (2017) Intra-neuronal Competition for Synaptic Partners Conserves the Amount of Dendritic Building Material. Neuron 93(3):632–645.

Savalli N, Pantazis A, Sigg D, Weiss JN, Neely A, Olcese R (2016) The α2δ-1 subunit remodels CaV1.2 voltage sensors and allows Ca2+ influx at physiological membrane potentials. J Gen Physiol 148(2):147–159.

Schmitt S, Evers JF, Duch C, Scholz M, Obermayer K (2004) New methods for the computer-assisted 3-D reconstruction of neurons from confocal image stacks. Neuroimage 23(4): 1283–1298.

Schützler N, Girwert C, Hügli I, Mohana G, Roignant JY, Ryglewski S, Duch C (2019) Tyramine action on motoneuron excitability and adaptable tyramine/octopamine ratios adjust Drosophila locomotion to nutritional state. Proc Natl Acad Sci U.S.A. 116(9):3805–3810.

Scott MB, Kammermeier PJ (2017) CaV2 channel subtype expression in rat sympathetic neurons is selectively regulated by α2δ subunits. Channels (Austin). 11(6):555–573.

Tong XJ, López-Soto EJ, Li L, Liu H, Nedelcu D, Lipscombe D, Hu Z, Kaplan JM (2017) Retrograde Synaptic Inhibition Is Mediated by α-Neurexin Binding to the α2δ Subunits of N-Type Calcium Channels. Neuron 95(2):326–340.

Venken KJ, Schulze KL, Haelterman NA, Pan H, He Y, Evans-Holm M, Carlson JW, Levis RW, Spradling AC, Hoskins RA, Bellen HJ (2011) MiMIC: a highly versatile transposon insertion resource for engineering Drosophila melanogaster genes. Nat Methods 8(9):737–743.

Vonhoff F, Duch C (2010) Tiling among stereotyped dendritic branches in an identified Drosophila motoneuron. J Comp Neurol 518(12):2169–2185.

Vonhoff F, Kuehn C, Blumenstock S, Sanyal S, Duch C (2013) Temporal coherency between receptor expression, neural activity and AP-1-dependent transcription regulates Drosophila motoneuron dendrite development. Development 140(3):606–616.

Wagh DA, Rasse TM, Asan E, Hofbauer A, Schwenkert I, Dürrbeck H, Buchner S, Dabauvalle MC, Schmidt M, Qin G, Wichmann C, Kittel R, Sigrist SJ, Buchner E (2006) Bruchpilot, a protein with homology to ELKS/CAST, is required for structural integrity and function of synaptic active zones in Drosophila. Neuron 49(6):833–844. Wagh DA, Rasse TM, Asan E, Hofbauer A, Schwenkert I, Dürrbeck H, Buchner S, Dabauvalle MC, Schmidt M, Qin G, Wichmann C, Kittel R, Sigrist SJ, Buchner E Bruchpilot, a protein with homology to ELKS/CAST, is required for structural integrity and function of synaptic active zones in Drosophila. Neuron 51(2):275 (2006).

Wang T, Jones RT, Whippen JM, Davis GW (2016) α2δ-3 Is Required for Rapid Transsynaptic Homeostatic Signaling. Cell Rep 16(11):2875–2888.

Worrell JW, Levine RB (2008) Characterization of voltage-dependent Ca2+ currents in identified Drosophila motoneurons in situ. J Neurophysiol 100(2):868–878.

